# A novel and evolutionarily distinct flavoprotein monooxygenase drives skatole degradation in *Rhodococcus*

**DOI:** 10.64898/2025.12.19.694838

**Authors:** Nan Meng, Qiao Ma

## Abstract

Skatole (3-methylindole), a persistent and toxic *N*-heterocyclic aromatic pollutant, poses significant bioremediation challenges due to poorly defined biodegradation processes. This study systematically investigated the genetic determinants and metabolic mechanisms of skatole degradation in the environmentally versatile Gram-positive genus *Rhodococcus*. We demonstrated that skatole degradation was a broadly conserved trait across diverse *Rhodococcus* species, including *R. aetherivorans*, *R. pyridinivorans*, *R. ruber* and *R. qingshengii*. Genomic and functional analyses uncovered a novel flavoprotein monooxygenase (FPMO) SkaA as the initial catalyst for skatole catabolism. Heterologous expression in *Escherichia coli* confirmed SkaA’s essential catalytic role, with high-resolution liquid chromatography-tandem mass spectrometry identifying the key product 3-methyloxindole and 3-hydroxy-3-methyloxindole. Distribution studies revealed SkaA homologs predominantly in Actinobacteria, particularly *Nocardia* and *Rhodococcus*, indicating conserved metabolic capacity for skatole transformation within this phylum. Notably, *Rhodococcus* strains lacking the *skaA* gene also retained skatole degradation activity, implying the existence of alternative genetic determinants. Crucially, phylogenetic analysis positioned SkaA as a distinct subclass within Group E FPMOs, exhibiting ≤40% sequence identity to reported styrene/indole oxygenases. The phylogenetic segregation of skatole, indole, and styrene monooxygenases provides a predictive framework for functional annotation of Group E FPMOs. Our findings elucidate a novel skatole transformation pathway, establish *Rhodococcus* as environmentally versatile biocatalysts, and provide new insights into the study of Group E FPMOs.

**IMPORTANCE:** Skatole is a notorious, foul-smelling compound generated by the anaerobic breakdown of tryptophan, presenting substantial risks to both the environment and public health. Despite its prevalence, the enzymatic machinery initiating its biodegradation remains poorly characterized. Our study resolves this critical gap by identifying SkaA as a functionally validated enzyme catalyzing skatole’s initial oxidation step. Distribution analyses revealed SkaA homologs predominantly in Actinobacteria, particularly *Rhodococcus* and *Nocardia*. Phylogenetic analysis positioned SkaA as a novel third subclass within Group E flavoprotein monooxygenases. These breakthroughs provide molecular tools for engineering targeted bioremediation solutions and new insights into the study of Group E FPMOs.

## INTROCUTION

Odorous emissions, comprising complex mixtures of inorganic (e.g., H₂S, NH₃) and organic pollutants (e.g., volatile fatty acids, cresols, indoles), pose significant challenges to environmental quality, animal welfare, and public health across livestock, aquaculture, landfill, and wastewater treatment sectors (1–4). Among these compounds, skatole (3-methylindole) is distinguished by its exceptional recalcitrance and malodor, attributable to its exceptionally low odor threshold (0.327 ng/L, as determined by expert panels), environmental persistence, and documented cytotoxicity (5). As a tryptophan-derived microbial metabolite, skatole accumulates to concerning levels (e.g., up to 20 mg/L in wastewater, 100 mg/kg in livestock manure) driven by anaerobic bacteria (6, 7). While recent advances have elucidated its biosynthetic pathways, primarily mediated by *Clostridium* and *Olsenella*, the environmental fate and transformation mechanisms of skatole remain poorly resolved, limiting the development of effective risk mitigation strategies (8, 9).

Microbial degradation represents a promising approach for skatole remediation, with diverse taxa reported to metabolize this compound. Current studies have focused predominantly on aerobic degradation, identifying several Gram-negative genera (*Pseudomonas*, *Acinetobacter*, *Cupriavidus*, *Rhodopseudomonas*, and *Burkholderia*) capable of skatole catabolism, with tentative pathways proposed (10–18). Conversely, Gram-positive degraders are less characterized, with reports largely restricted to *Rhodococcus* strains. *Rhodococcus* spp. exhibit superior environmental resilience, broad substrate spectra, and redundant metabolic pathways for aromatic pollutants (19–24). Their capacity to colonize diverse niches (e.g., soil, sludge) further positions this genus as an ideal candidate for bioremediation applications. Our previous metagenomic and metatranscriptomic analyses identified *Rhodococcus* as a primary active population responsible for skatole degradation in activated sludge systems (20, 22). We further demonstrated the skatole degradation capacity and efficacy in strain *R. aetherivorans* DMU1 (20, 22). Subsequent studies have since confirmed degradation capabilities in other strains, including *R. aetherivorans* BCP1 and *R. ruber* R1 (24). These collective advances prompt investigation into whether skatole biodegradation constitutes a conserved catabolic trait within the *Rhodococcus* genus.

Despite progress in skatole-degrading strain isolation and characterization, critical knowledge gaps persist. First, the genetic basis for skatole degradation remains unconfirmed. Our pioneering work on *R. aetherivorans* DMU1 identified a potential oxidoreductase gene cluster involved in skatole conversion via RNA-seq and quantitative PCR analyses (20, 22). This cluster included a putative flavoprotein monooxygenase (FPMO), representing a distinct genetic module potentially dedicated to skatole biodegradation. While this finding has garnered recognition, the enzymatic activity, phylogenetic distribution, and uniqueness of this system remain unclear. Second, the biotransformation pathway of skatole is ambiguous. Although several intermediates have been tentatively identified via mass spectrometry, their structural assignments remain ambiguous, and the enzymatic catalytic steps are undefined (20, 22).

In this study, we systematically investigated skatole degradation in representative strains of the environmentally versatile *Rhodococcus* strains isolated in our laboratory. We tried to address three key questions: (I) Is skatole degradation a common trait among *Rhodococcus* strains? (II) What is the metabolic pathway for skatole degradation in *Rhodococcus* strains? and (III) What is the core functional gene responsible for skatole degradation, and how are they distributed? Our findings should provide new insights into the molecular mechanisms of skatole metabolism in Gram-positive bacteria and identify novel biocatalytic resources for environmental remediation.

## MATERIALS AND METHODS

### Chemicals and media

Skatole was purchased from Aladdin (Shanghai, China). All other chemicals and solvents were reagent grade and purchased from commercial sources. The culture media were prepared as follows. Mineral salt (MS) medium (per liter): (NH₄)₂SO₄ 134 mg, KH₂PO₄ 141 mg, K₂HPO₄ 287 mg, MnSO₄·H₂O 2.68 mg, MgSO₄·7H₂O 21.4 mg, FeCl₃·6H₂O 0.000134 mg, and CaCl₂ 3.8 mg. Mineral salt yeast extract (MSY) medium was prepared from MS medium by adding 1g/L yeast extract. Luria-Bertani (LB) medium (per liter): tryptone 10 g, NaCl 10 g, and yeast extract 5 g.

### Skatole degradation by *Rhodococcus* strains

*Rhodococcus* strains DMU1, DMU2, DMU114, DMU2021, E7, SJ-1, SJ-2, and SJ-3, which were isolated and preserved in our lab, were separately cultured in MSY medium. Upon reaching the logarithmic growth phase, cells were harvested, washed, and resuspended in phosphate-buffered saline (PBS, pH 7.0) to an OD₆₀₀ of 1.0. A 1% (*v*/*v*) inoculum was then transferred to both MS and MSY media supplemented with 40 mg/L skatole to evaluate degradation capability, with uninoculated media serving as controls. All groups included three biological replicates and were incubated at 30 °C with shaking at 150 rpm. Skatole degradation was monitored by sampling at 0 h, 48 h, and 96 h, followed by analysis using ultra-performance liquid chromatography (UPLC, Waters) under specified conditions: detection wavelength 280 nm, flow rate 0.8 mL/min, and mobile phase methanol:water (60:40, *v/v*).

### Genome analyses and strain identification

Draft genomes of strains DMU2 and E7 were generated using Illumina sequencing, and the genomes of the other six strains were obtained previously (22, 25, 26). Comparative genomic analysis of all eight *Rhodococcus* strains was performed with BRIG 1.4.1 software, utilizing the complete genome of *R. aetherivorans* DMU1 as the reference. For taxonomic classification, pairwise genome comparisons against 45 validated *Rhodococcus* species were conducted through Average Nucleotide Identity (ANI) analysis using the JSpeciesWS web service (https://jspecies.ribohost.com/jspeciesws/) (27). Digital DNA-DNA hybridization (dDDH) values derived from the d4 formula were calculated via the TYGS platform (https://tygs.dsmz.de/) (28). With the criteria of average nucleotide identity (ANI) ≥ 95% and dDDH ≥ 70% for species delineation, the taxonomic species of the eight strains were determined.

### Quantitative real-time RT-PCR (qPCR) assay

Gene expression was analyzed by quantitative real-time PCR (qPCR). Total RNA was extracted from bacterial strains using a Vazyme RNA extraction kit and reverse-transcribed into cDNA using the Hiscript III Reverse Transcriptase kit (Vazyme, China). The qPCR was performed using ChamQ Universal SYBR qPCR Master Mix (Vazyme, China) on a real-time PCR system (LightCycler 480II, Roche, Switzerland). The 16S rRNA gene was used as the internal reference for normalization. The relative expression level of the target gene was calculated using the 2^-ΔΔCT^ method. All reactions were performed in triplicate and primers were listed in Table S1.

### Gene expression and function verification

Primers targeting *skaA* from strain DMU1 and SJ-1 (Table S1) were designed for directional cloning into pET-28a(+) expression vector. Gene *skaA* from strain R1 was synthesized by Sangon Biotech (Shanghai, China) and expressed in pET-32a(+) using NcoI and XhoI cloning sites. The resultant recombinant plasmid was transformed into *E. coli* DH5α for sequence verification before electroporation into *E. coli* BL21(DE3) for functional studies. Transformants were cultivated in Luria-Bertani (LB) medium with 100 mg/L kanamycin or 50 mg/L ampicillin at 37°C and 150 rpm. The *skaA* heterologous expression strain was induced at an OD_600_ of 0.4 by adding 0.2 mM isopropyl-β-d-thiogalactoside and then incubated for 20 h at 30°C. For resting cell assays, harvested cells were washed and resuspended in PBS. Reactions initiated with 50 mg/L skatole (treatment) or PBS alone (control) were incubated at 30°C with shaking.

### Metabolite identification

Skatole transformation metabolites of strains DMU1 and *E. coli* BL21 expressing *skaA* were analyzed by chromatography–high-resolution mass spectrometry (LC-HRMS). Analyses were performed on a Thermo Fisher Q Exactive Plus Orbitrap mass spectrometer equipped with an electrospray ionization source in positive ion mode. The spray voltage was 3.6 kV, capillary temperature 320 °C, ion source temperature 320 °C, sheath gas flow 35, auxiliary gas flow 10, and probe heater (desolvation line) temperature 450 °C. Chromatographic separation was carried out on a Thermo Scientific Hypersil GOLD C18 column (100 × 2.1 mm, 1.9 µm) using the mobile phase of 0.1% formic acid in water (A) and acetonitrile (B). A gradient elution was applied as follows: 0–2 min, 5% B; 2–13 min, 5–100% B; 13–16 min, 100% B; 16–20 min, 5% B, at a flow rate 0.2 mL/min and column temperature 40 °C.

### Distribution analysis of SkaA homologs

To investigate the distribution of skatole oxygenase homologs across bacterial genomes, we screened two comprehensive microbial proteome databases constructed from NCBI resources: the RefSeq database (333,218,737 predicted protein sequences, March 2025) and the non-redundant (NR) database (707,028,945 predicted protein sequences, April 2025) (29). Using DIAMOND v2.1.11 with uniform parameters (sequence identity >40%, query/subject coverage >70%), we queried both databases against the reference skatole oxygenase SkaA (UniProt: QIX49842) from *R. aetherivorans* DMU1. Homologous sequences were taxonomically classified to determine phylum- and genus-level distribution patterns. For phylogenetic reconstruction, candidate homologs were aligned with MAFFT (default settings), followed by maximum-likelihood tree inference using FastTree v2.1.11. Evolutionary relationships were visualized and annotated in iTOL.

### Analysis of the *ska* gene cluster in skatole-degrading strains

The SkaA protein sequence from strain DMU1 served as the reference for BLASTP analysis (E-value ≤1×10⁻⁵) against the proteomes of all reported skatole-degrading bacteria retrieved from NCBI. Matching sequences were mapped to corresponding coding sequences (CDS) in GenBank annotations to identify homologous monooxygenase gene clusters. Gene cluster visualizations were generated using ChiPlot (https://www.chiplot.online/).

### Phylogenetic analysis of skatole, indole, and styrene monooxygenases

Phylogenetic analysis of the functionally validated skatole, indole, and styrene monooxygenases was conducted by aligning the protein sequences using ClustalW. The aligned sequences were used to construct a Maximum Likelihood (ML) phylogenetic tree with 1000 bootstrap replicates under the Jones-Taylor-Thornton amino acid substitution model by MEGA12 (30).

## RESULTS

### Skatole degradation is prevalent in *Rhodococcus* strains

Skatole degradation has been predominantly investigated in Gram-negative bacteria, including *Pseudomonas*, *Acinetobacter*, *Burkholderia*, and *Cupriavidus* (Table S2). *Rhodococcus*, a genus of typical Gram-positive bacteria, possesses large genomes with numerous redundant aromatic-degrading genes, making its strains promising microbial resources for bioremediation (31). Notably, recent studies have identified several *Rhodococcus* species as highly efficient skatole degraders, including *R. aetherivorans* DMU1, *R. pyridinivorans* Rp3, and *R. ruber* R1 (21, 23, 22, 24). These findings motivated us to assess whether skatole degradation capability is a prevalent trait within the *Rhodococcus* genus.

The skatole-degrading capacity of eight *Rhodococcus* strains, previously isolated in our laboratory from activated sludge and marine sediment samples, was assessed (19). Degradation performance was evaluated using two culture media: MS (skatole as the sole carbon source) and MSY (supplemented with yeast extract as an additional carbon source) (Table 1). In MS medium, strains DMU1, DMU2, and SJ-2 demonstrated the highest degradation efficiency, achieving complete skatole removal within 48 hours (Fig. S1). Strains DMU114 and DMU2021 required 96 hours for >99% degradation. Conversely, strains E7 (27.7%, 96 hours), SJ-1 (34.0%), and SJ-3 (6.1%) exhibited significantly weaker degradation activity. In MSY medium, strains DMU1, DMU114, SJ-1, SJ-2, and SJ-3 degraded >99% of skatole within 96 hours (Fig. S2). However, the degradation efficiencies of strains DMU2, DMU2021, and SJ-2 were lower in MSY compared to MS medium, suggesting potential catabolite repression in the presence of yeast extract. Conversely, a particularly significant metabolic shift was observed in strains SJ-1 and SJ-3. They degraded skatole inefficiently in MS but achieved complete removal within 48 hours in MSY. Strain DMU114 also demonstrated faster degradation in MSY medium (77.5% degradation at 48 hours) compared to MS medium (11.0%, 48 hours). This suggested that the presence of yeast extract in MSY medium enabled co-metabolism of skatole. Among the tested strains, DMU1 exhibited superior degradation performance, whereas strain E7 displayed the least activity.

**Table 1.** Skatole-degrading abilities of the *Rhodococcus* strains

| Strain | MS-48 h | MS-96 h | MSY-48 h | MSY-96 h |
| --- | --- | --- | --- | --- |
| DMU1 | 100% | 100% | 97.7% | 100% |
| DMU2 | 100% | 100% | 71.2% | 80.5% |
| DMU114 | 11.0% | 100% | 77.5% | 100% |
| DMU2021 | 5.4% | 99.2% | 34.2% | 53.7% |
| E7 | 19.5% | 27.7% | 24.2% | 32.5% |
| SJ-1 | 12.2% | 34.0% | 100% | 100% |
| SJ-2 | 100% | 100% | 76.8% | 99.2% |
| SJ-3 | 3.7% | 6.1% | 100% | 100% |

Collectively, all evaluated *Rhodococcus* strains demonstrated the capacity to degrade skatole, albeit with performance being strongly strain-specific and medium-dependent. These phenotypic observations led us to hypothesize the potential presence of conserved genetic determinants that enabled skatole degradation across the genus. To identify these mechanisms, whole-genome sequencing and comparative analysis of the strain genomes were performed.

### Genome sequencing and classification of *Rhodococcus* strains

The complete genome sequences of strains DMU1, DMU114, DMU2021, SJ-1, SJ-2, and SJ-3 were available from our previous studies (20, 26, 32, 33), and draft genome sequences of strains E7 and DMU2 were generated in this work. Genome information for all eight strains is summarized in Table 2. A comparative genomic analysis was performed using a circular map, employing the genome of strain DMU1 as a reference. Genes were color-coded based on their similarity to orthologs in DMU1 (Fig. S3). Genomes ranged in size from 5.2 to 6.8 Mb, contained 5219–6498 coding sequences, and possessed GC contents of 62.5–70.0%.

**Table 2.** Genome information of the *Rhodococcus* strains used in this study

| Strain | Genus | Species | Source | GC content | Genome size | Coding sequences |
| --- | --- | --- | --- | --- | --- | --- |
| DMU1 | *Rhodococcus* | *aetherivorans* | Activated sludge | 70.0 | 6835256 | 6219 |
| DMU2 | *Rhodococcus* | *qingshengii* | Activated sludge | 62.5 | 6634658 | 6498 |
| DMU114 | *Rhodococcus* | *pyridinivorans* | Activated sludge | 67.9 | 5241731 | 5219 |
| DMU2021 | *Rhodococcus* | *pyridinivorans* | Activated sludge | 67.7 | 5511496 | 5439 |
| E7 | *Rhodococcus* | *qingshengii* | Marine sediment | 62.5 | 6289418 | 6042 |
| SJ-1 | *Rhodococcus* | *ruber* | Marine sediment | 70.4 | 5738084 | 5530 |
| SJ-2 | *Rhodococcus* | unknown | Marine sediment | 65.4 | 5773915 | 5878 |
| SJ-3 | *Rhodococcus* | unknown | Marine sediment | 65.4 | 5833229 | 5692 |

Taxonomic classification of all eight strains was determined through whole-genome ANI and dDDH analyses (Tables S3 and S4). ANI values between the experimental strains and 45 representative *Rhodococcus* strains ranged from 69.6% to 98.8% (Fig. S4). Pairwise comparisons revealed that *R. aetherivorans* JCM 14343 exhibited the highest ANI (97.2%) and dDDH (91.4%) values with strain DMU1, confirming its species affiliation. Strains DMU2 and E7 showed the highest ANI similarity to *R. qingshengii* JCM 15477 (98.4% and 98.6%). The dDDH values between *R. qingshengii* JCM 15477 and DMU2 and E7 were 88.5% and 69.5%, respectively. Strains DMU114 and DMU2021 displayed the highest mutual ANI similarity (98.1%). DMU2021 exhibited its highest ANI value (98.4%) and dDDH (91.1%) with *R. pyridinivorans* DSM 44555, identifying its species affiliation. Strain SJ-1 demonstrated the highest ANI similarity (98.8%) and dDDH (92.7%) with *R. ruber* DSM 43338, confirming classification within this species. Strains SJ-2 and SJ-3 exhibited the highest ANI similarity to each other (>98%), whereas ANI values against all other investigated *Rhodococcus* species were below 95% and dDDH values were below 35%, indicating these might belong to a novel species which required further verification.

Based on these genomic analyses, strains were assigned as follows. DMU1: *R. aetherivorans*; DMU2 and E7: *R. qingshengii*; DMU114 and DMU2021: *R. pyridinivorans*; SJ-1: *R. ruber*; while SJ-2 and SJ-3 were unknown species (Table 2). Three species, i.e., *R. aetherivorans* (strain DMU1), *R. pyridinivorans* (strain Rp3), and *R. ruber* (strain R1), have been reported to catalyze skatole (19, 20, 22). Our study further established the identified *Rhodococcus* strains, including the validated *R. qingshengii* and a potentially novel species, as new microbial resources for skatole biodegradation. Notably, despite both being classified as *R. qingshengii*, strains E7 (<33% degradation in MS/MSY) and DMU2 (complete degradation within 48 h in MS medium) exhibited profoundly different skatole degradation efficiencies, further confirming the critical role of intra-species variation in degradation performance (Table 1).

### Functional verification of the skatole oxygenase gene in strain DMU1

In a previous study, a gene *skaA* (1368 bp) encoding a FPMO was identified in *R. aetherivorans* DMU1 and implicated as the key enzyme in skatole degradation by transcriptomic analysis (22). qPCR analysis confirmed that gene *skaA* was significantly upregulated in response to both skatole and indole, suggesting its functional involvement in the degradation of both compounds (Fig. S5). The *skaA* gene was successfully cloned and expressed in in *E. coli* BL21(DE3) strain (Fig. S6). Notably, this engineered strain was capable of skatole biotransformation and could also convert indole into indigoid pigments (Fig. S6).

### Skatole transformation product identification

Metabolite analysis was performed using resting cells, with reaction mixtures subsequently analyzed by LC-HRMS. Three major products, each exhibiting an [M+H]⁺ ion at *m*/*z* 148.07569, were detected in positive ion mode with retention time (RT) of 5.37, 7.82, and 9.66 min. These compounds shared the molecular formula C₉H₉NO and yielded identical MS/MS fragmentation patterns, indicating they were structural isomers derived from the monooxygenation of skatole. Product I (RT 7.82 min) was identified as 3-methyloxindole by comparing it to an authentic standard (Fig. 1a, Fig. S7a). The products eluting at 5.37 min and 9.66 min were tentatively proposed as 3-methyl-3H-indol-3-ol and 2,3-epoxy-3-methylindole, a characteristic epoxide intermediate formed by Group E FPMOs(34). Additionally, product II (RT 6.24 min, *m*/*z* 146.06018 [M+H]⁺) was confirmed as 3-hydroxy-3-methyloxindole by comparison with a reference standard (Fig. 1b, Fig. S7b).

**Fig. 1.**
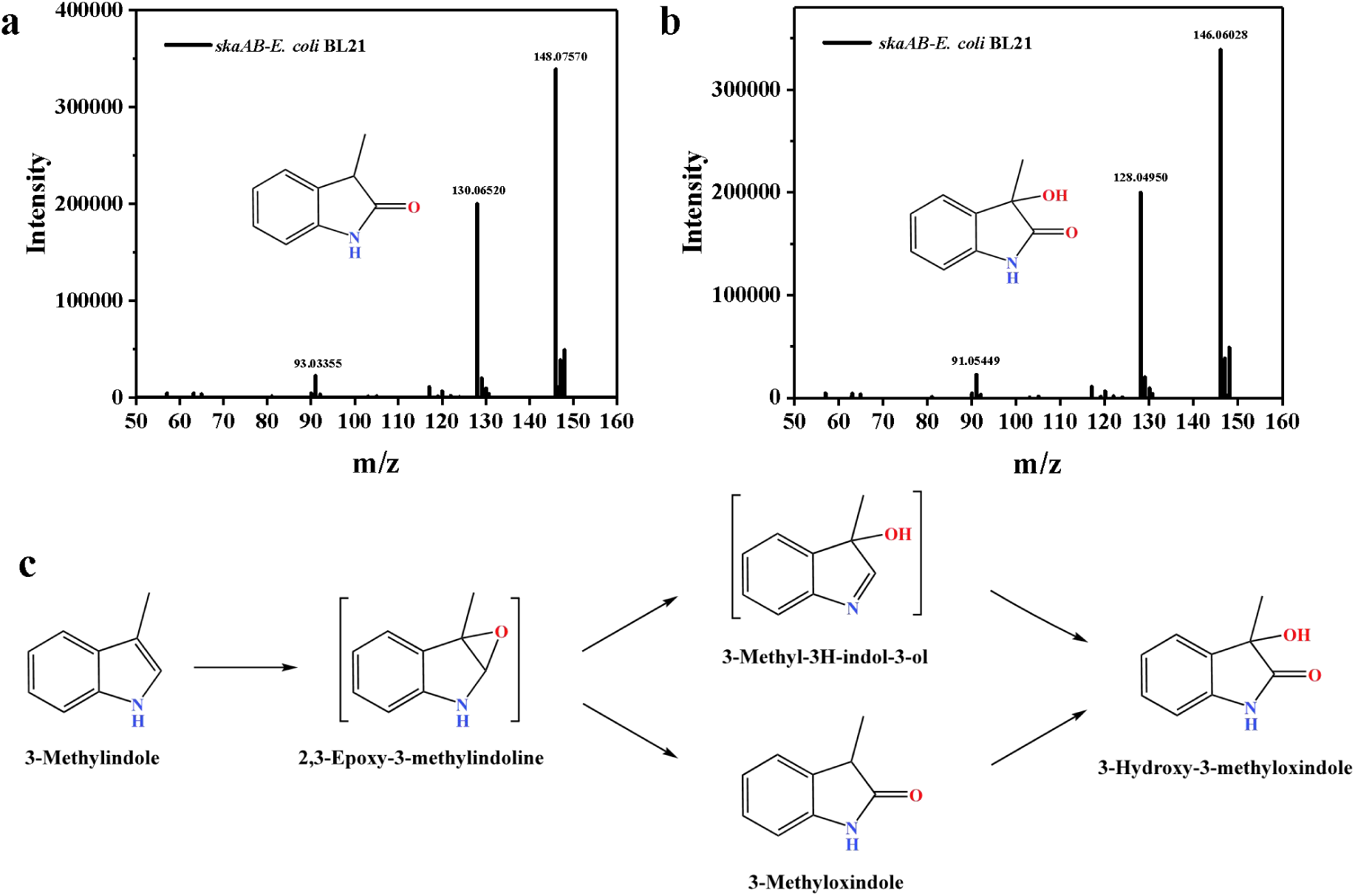
Identification of the skatole transformation metabolites in gene engineering *E. coli* BL21 (DE3)-*skaAB* strain. (a) Mass spectrum of product 3-methyloxindole. (b) Mass spectrum of 3-hydroxy-3-methyloxindole. (c) Proposed skatole transformation pathway.

Collectively, these results suggested a novel skatole biotransformation pathway (Fig. 1c). Skatole was initially oxygenated by SkaA to form the epoxide intermediate 2,3-epoxy-3-methylindole. This unstable species was proposed to undergo subsequent further transformations to yield 3-methyloxindole (RT 7.82 min), which was further converted to 3-hydroxy-3-methyloxindole. Analysis of wild-type strain *R. aetherivorans* DMU1 metabolites also identified both 3-methyloxindole (*m*/*z* 148.07569, RT 7.85 min) and 3-hydroxy-3-methyloxindole (*m*/*z* 146.04006, RT 6.44 min), confirming their relevance in the native metabolic pathway.

### Skatole monooxygenase SkaA is not the sole enzyme involved in skatole transformation

Genomic analysis of the *skaA* gene across all *Rhodococcus* strains used in this study revealed its presence in multiple species (Table S5). Putative skatole monooxygenases (SkaMOs) exhibiting varying degrees of protein sequence identity (>30%) were identified in *R. aetherivorans* JCM 14343 (38.2% and 99.3%), *R. indonesiensis* SARSHI1 (38.4%), *R. opacus* ATCC 51881 (30.7%, 33.4%, and 37.9%), *R. sacchari* Z13 (36.9%), *R. spongiicola* LHW 50502 (31.9% and 34.0%), and *R. triatomae* DSM 44893 (36.6%). Notably, SkaMO-like gene was also detected in *R. ruber* SJ-1 (38.2%). However, similar sequences did not exist in strains DMU2, DMU114, DMU2021, E7, SJ-2, and SJ-3. Strikingly, pairwise comparison showed the SkaA sequence from strain DMU1 shares low identity (<40%) with orthologs in all listed strains except its conspecific strain *R. aetherivorans* JCM 14343, indicating conservation primarily within this species.

Furthermore, analysis of all currently available skatole-degrader genomes (Table S6) revealed putative SkaMOs in *R. ruber* R1 (38.2% and 98.9%), *R. aetherivorans* BCP1 (38.2% and 99.1%), *Cupriavidus* sp. KK10 (32.8% and 34.8%), *Burkholderia* sp. IDO3 (32.2% and 35.7%), and *Acinetobacter piscicola* p38 (31.6% and 36.9%). Notably, the recently characterized degrader *R. pyridinivorans* Rp3 lacked detectable SkaMOs, a finding consistent with strains *R. pyridinivorans* DMU2021 and DMU114, indicating novel degradation mechanisms within this species.

Comparative analysis of the *ska* gene clusters across *Rhodococcus* strains revealed a conserved functionally modular architecture in key degraders, including strains DMU1, R1, and BCP1 (20, 24). As illustrated in Fig. 2, this 14-gene cluster (*skaCDZEFGHAIJKRLB*) was transcriptionally upregulated under skatole stress in both strains DMU1 and R1. Bioinformatic analysis indicated that the initial oxidation could be mediated by the *skaAB*-encoded monooxygenase-reductase complex catalyzing skatole epoxidation. Gene *skaB* encoded a flavin reductase, which should provide the reducing equivalents to the oxygenase SkaA. A *skaB* gene (531 bp) was identified 7058 bp downstream of *skaA*, rather than in an adjacent position. Subsequent ring cleavage was likely executed by a three-component aromatic ring-hydroxylating oxygenase (ARHO) system (*skaCDFG*), while cluster expression was potentially governed by the embedded LuxR-type regulator *skaR* Table S7). Strain DMU1 possessed a single genomic *skaA* copy. However, strains R1 and BCP1 harbored an additional SkaA homolog exhibiting low sequence similarity to SkaA_DMU1 (38.2%), with corresponding truncated clusters containing only *skaAB* genes. This gene cluster was similar with that of strain SJ-1 (Fig. 2). The specific roles of the two gene clusters in the degradation of skatole require further validation.

**Fig. 2.**
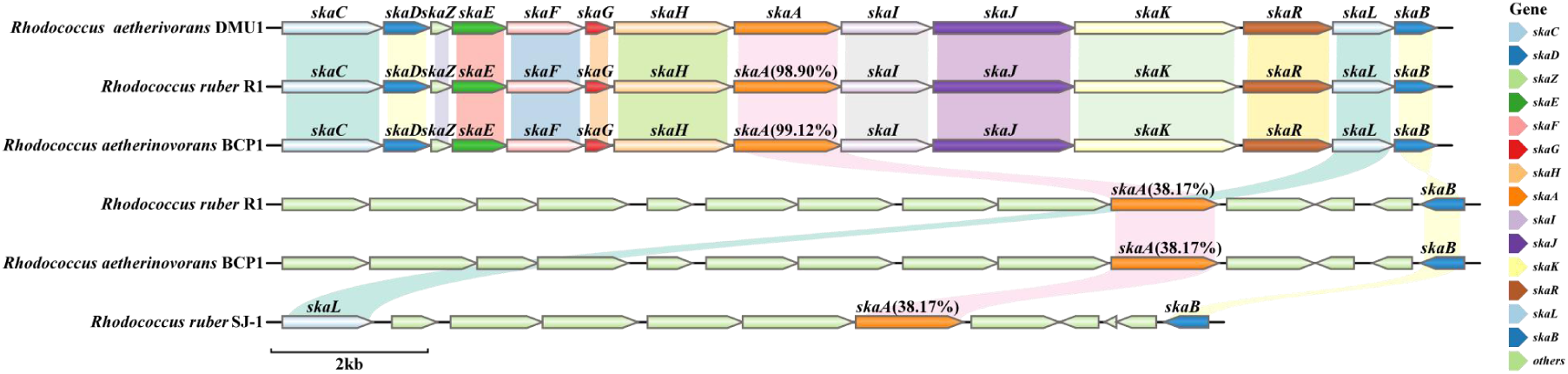
Gene cluster analysis of typical skatole-degrading *Rhodococcus* strains. Accession numbers for the strain DMU1 proteins are in Table S7, with the IDs of SkaA homologs from other strains provided in Table S6.

### Taxonomic distribution of SkaA in RefSeq and NR Databases

Sequence similarity analysis using the NCBI RefSeq database (to date March 2025; 36754 genomes, 333218737 proteins) identified 1492 putative SkaA homologs (≥30% amino acid identity). However, only 39 homologs exhibiting≥40% identity were identified (Table S8). These were distributed across 10 actinobacterial genera (*Antrihabitans*, *Arthrobacter*, *Cryobacterium*, *Gordonia*, *Haliangium*, *Nocardia*, *Pseudarthrobacter*, *Rhodococcus*, *Tsukamurella*, Vitiosangium) within the phylum Actinomycetota (Fig. 3). *Nocardia* (20 sequences, 51.3%) and *Rhodococcus* (9 sequences, 23.1%) were the predominant genera. This taxonomic distribution aligned with the established role of Actinomycetota in aerobic aromatic catabolism, particularly via oxygenase-dependent pathways. The prevalence of *Nocardia* and *Rhodococcus*, genera renowned for diverse redox enzyme systems, suggested they may be key contributors to environmental skatole degradation. While *Rhodococcus* skatole degraders have been recently documented, the potential role of *Nocardia* warrants investigation. It was notable that SkaA sequences within *R. aetherivorans* exhibited >99% identity, indicating high conservation (24). SkaA homolog with 98.9% identity was also identified in *R. ruber*, consistent with its recently documented skatole-degrading capability. In contrast, the *Nocardia* homologs displayed lower similarity (40.3–42.0%).

**Fig. 3.**
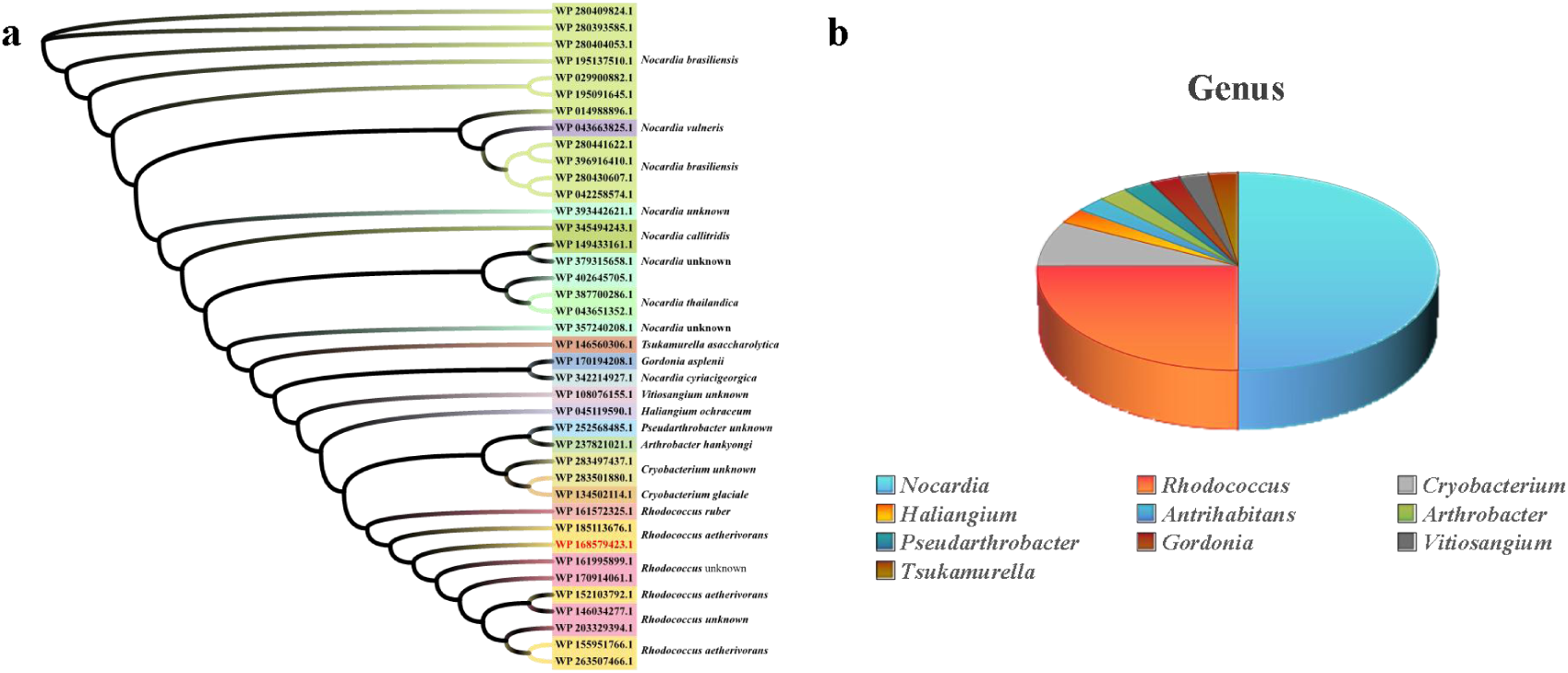
Distribution of SkaA homologs in related strains in the RefSeq database. (a) Phylogenetic tree for 40% identity in skate monooxygenases. (b) Genus distribution

Analysis of the NCBI non-redundant (NR) database (to date April 2025) expanded the dataset to 56 homologs (≥40% identity), spanning 19 genera across four phyla: Actinomycetota (41, 73.2%), Myxococcota (9, 16.1%), Acidobacteriota (4, 7.1%), and Pseudomonadota (2, 3.6%) (Fig. S9, Table S9). *Nocardia* (15 sequences) and *Rhodococcus* (14 sequences) remained dominant. This broader analysis revealed novel homologs in understudied phyla, particularly Myxococcota.

### Skatole monooxygenase belongs to a new branch of Group E FPMOs

FPMOs constitute a versatile class of biocatalysts that mediate chemo-, regio-, and enantioselective oxyfunctionalization reactions across diverse biological processes (35). Based on conserved sequence and structural features, FPMOs are categorized into eight distinct phylogenetic groups (group A-H). To resolve their evolutionary relationships and domain architectures, we constructed a comprehensive phylogenetic tree using representative FPMO sequences coupled with domain analysis (Fig. 4, Table S10). Group A universally contains the Pfam01494 domain (FAD_binding_3 superfamily cl21454), mediating flavin adenine dinucleotide (FAD) binding(36). Group B, also a member of the cl21454 superfamily, is characterized by the NAD_binding_8 and Pyr_redox_3 domains for NAD(P)H binding and pyridine ring oxidation-reduction reactions, respectively. Group C features the luciferase-like Pfam00296 domain, utilizing reduced flavin cofactors (FMNH₂/FADH₂) for oxygen activation rather than direct flavin substrate binding (37). Group D is characterized by dual domains 4-hydroxyphenylacetate 3-hydroxylase C-terminal HpaB (Pfam03241) and HpaB_N (Pfam11794) in hydroxylation cascades. Group E is defined by conserved styrene monooxygenase domains catalyzing epoxidation. Group F comprises tryptophan halogenases mediating regioselective halogenation. Group G contains Pfam01593 (flavin-dependent amine oxidoreductases) with integrated NAD_binding domains (38). Group H belongs to the cl21457 superfamily and possesses TIM-like beta/alpha barrel domains.

**Fig. 4.**
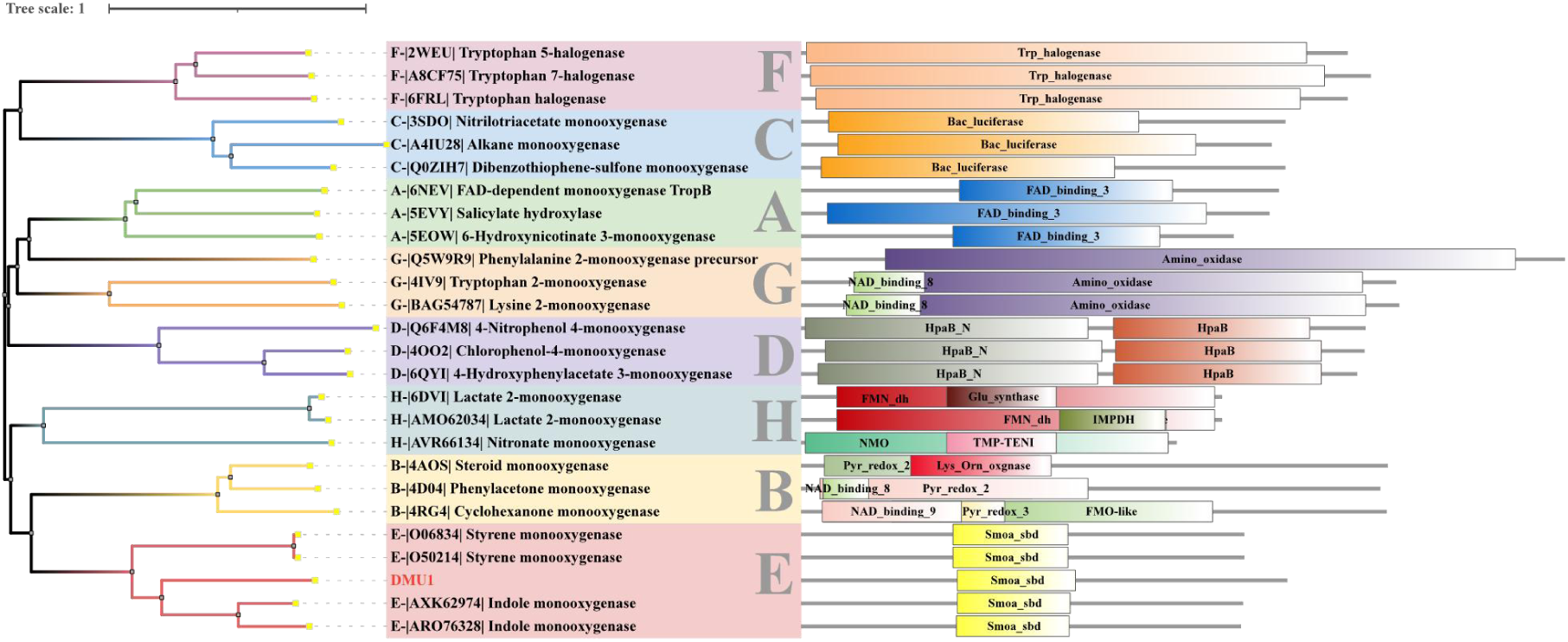
Phylogenetic and domain analyses of different types of FPMOs. Related information was provided in Table S10.

Phylogenetic analysis unequivocally positioned the SkaMO within Group E FPMOs (Fig. 4). In this family, a flavin reductase provided the monooxygenase subunit with reduced FAD (34). Group E FPMOs were primarily divided into styrene monooxygenases (SMOs) and indole monooxygenases (IMOs) based on substrate specificity (30). Comparative sequence analysis revealed SkaA-DMU1 shared limited homology with established subtypes: 23.8-29.0% identity with SMOs and 27.7-35.1% with IMOs (Table S11). Despite showing closer affinity to IMOs, SkaMOs lacked conserved catalytic motifs characteristic of both subtypes - specifically the SMO fingerprint (N46-V48-H50-Y73-H76-S96) and IMO fingerprint (S46-Q48-M50-V/I73-I76-A96) (39).

Comprehensive phylogenetic reconstruction including all characterized IMOs/SMOs demonstrated three evolutionarily divergent branches: IMOs, SMOs (including SMO-like enzymes with unknown physiological role), and SkaMOs (Fig. 5). Remarkably, the SkaMOs clade formed a monophyletic cluster comprising both Gram-positive (*Rhodococcus* spp.) and Gram-negative (*Burkholderia*, *Acinetobacter*) variants despite low sequence identity (<40%) (22, 24, 21, 40, 41, 12, 15). This phylogeny enabled functional predictions for several oxygenases: (I) Previously reported *Gordonia rubripertincta* IndA (ASR05096) likely represented a SkaMO homolog (42, 30); (II) The IifC1 (APT36898) from strain *Burkholderia* sp. IDO3 might be a SkaMO while IifC2 (AXK62947) was an IMO (43, 40). (III) Protein XEZ56878 from strain *Acinetobacter piscicola* p38 was a SkaMO, while XEZ56851 was an IMO (15). These findings established SkaMO as a mechanistically distinct subclass within Group E FPMOs, necessitating expansion of current classification frameworks.

**Fig. 5.**
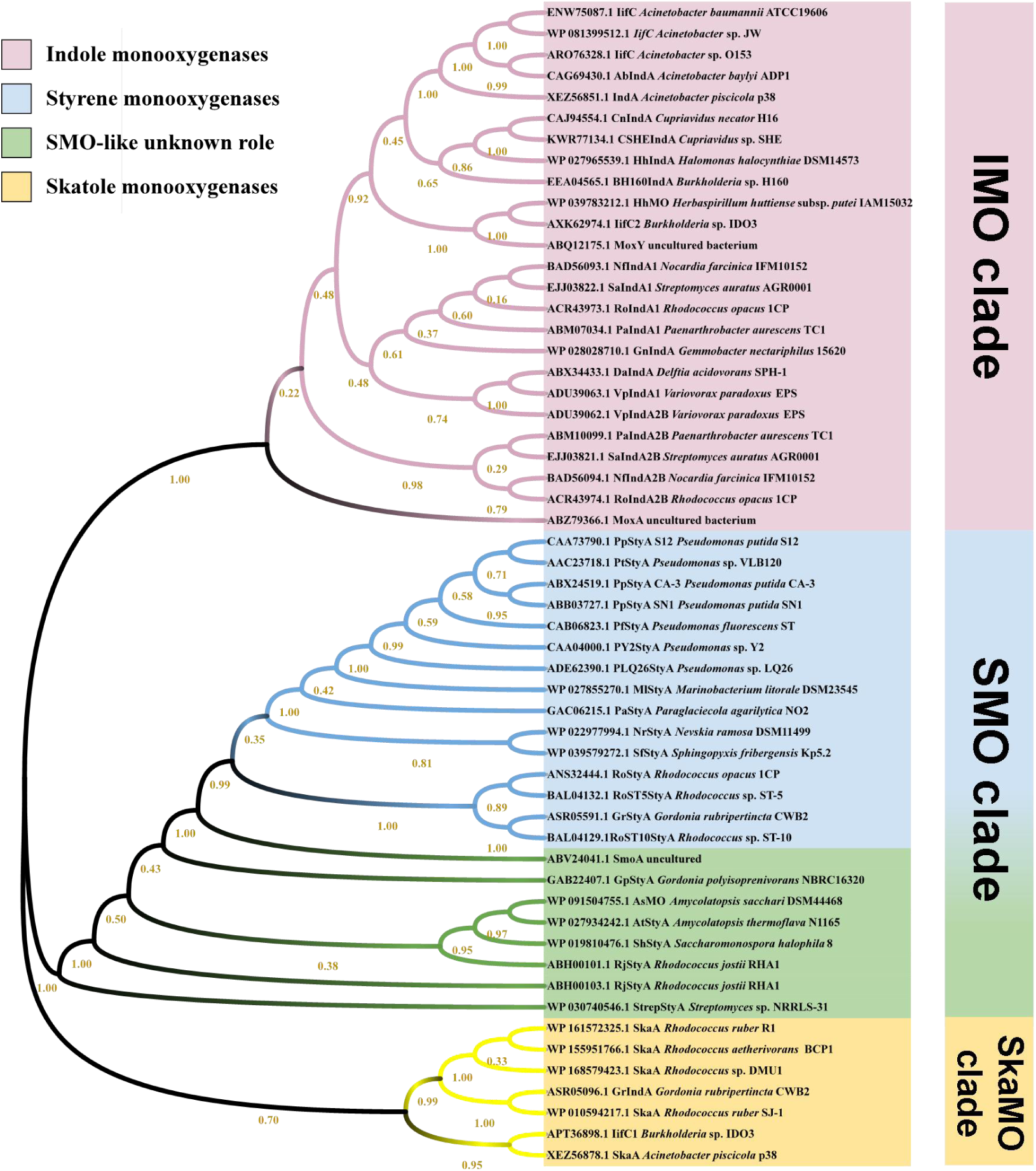
Phylogenetic tree analysis of the reported Group E FPMOs. IMO, indole monooxygenase, in purple. SMO (styrene monooxygenase) with know roles, in blue. SMO-like enzymes with unknown physiological role, in green. SkaMO, skatole monooxygenase, in yellow.

To further verify the funcions of SkaA homologs in skatole transformtion, we expressed the SJ-1_*skaA*, and R1_*skaA* genes (Fig. S10a). The functional activity of heterologously expressed SkaA protein was evaluated by measuring skatole concentration over time in a resting bacterial cell system. It was shown that both R1-SkaA and SJ-1-SkaA displayed skatole degradation activity.

## DISCUSSION

The present study systematically deciphered the degradation performance, metabolic pathways, and genetic determinants within the environmentally versatile genus *Rhodococcus*.

We demonstrate that efficient skatole degradation occurs in eight different *Rhodococcus* strains and identify a novel FPMO SkaA as the initial catalyst for skatole oxidation. Crucially, phylogenetic analysis reveals SkaMOs define a distinct subclass within Group E FPMOs, diverging from well-documented IMOs and SMOs. These findings provide mechanistic insights into Gram-positive skatole catabolism and establish *Rhodococcus* as a promising biocatalyst for skatole remediation.

Our study firstly establishes skatole degradation as a conserved trait across the genus *Rhodococcus*. All eight tested strains, spanning different species, exhibited skatole degradation activity (Fig. S1, Fig. S2, Table 1). This significantly expands prior knowledge where only three species (*R. aetherivorans*, *R. ruber*, and *R. pyridinivorans*) were documented skatole degraders (44, 20, 22, 24, 23). While degradation efficiency is strain- and medium-dependent (e.g., *R. qingshengii* DMU2 achieved complete degradation in 48 h in minimal medium, whereas conspecific strain E7 showed limited activity), the universal capability across phylogenetically distinct isolates underscores *Rhodococcus*’s intrinsic adaptability as a skatole degrader. This aligns with the genus’s recognized genomic plasticity and redundant aromatic metabolic pathways (45, 46). Crucially, we identify *skaA* as a core functional gene conserved in *R. aetherivorans* and *R. ruber* (Fig. 5; Table S8), directly explaining their high degradation efficacy. Strikingly, *skaA*-independent degradation occurred in *R. pyridinivorans* (Rp3, DMU114, and DMU2021), *R. qingshengii* (E7, DMU2), and marine isolates SJ-2/SJ-3, demonstrating alternative genetic pathways and highlighting the genus’s metabolic versatility (47). The metabolic basis of skatole degradation in *skaA*-deficient strains remains to be elucidated, as the yet-to-be-identified pathway may represent another conserved mechanism across diverse *Rhodococcus* lineages (48). Critically, this functional versatility positions *Rhodococcus* as an effective biocatalyst for skatole deodorization, particularly in high-load environments like livestock manure systems We secondly elucidate a novel skatole epoxidation pathway initiated by SkaA, resolving key uncertainties in Gram-positive catabolism. LC-HRMS unambiguously identified 3-methyloxindole and 3-hydroxy-3-methyloxindole as metabolites in both engineered *E. coli* and wild-type *R. aetherivorans* DMU1 strain (Fig. 1). Consistent with well-established mechanisms for SMOs and IMOs, which epoxidize substrates to styrene oxide and indole-2,3-epoxide, we reasonably propose that SkaA directly generates 2,3-epoxy-3-methylindole (Fig. S8) (35, 30, 34). This unstable epoxide spontaneously rearranges to 3-methyl-3H-indol-3-ol and 3-methyloxindole—a metabolite previously detected in anaerobic microbial systems (e.g., methanogenic and sulfate-reducing conditions) and proposed in *Pseudomonas putida* LPC24, *Acinetobacter oleivorans* AO-06, and *Acinetobacter xiamenensis* Ya (11, 13, 14). 3-Hydroxy-3-methyloxindole formation from skatole was reported in an Gram-positive bacterium by Fujioka and Wada by an characterized enzyme (49). Our work confirms SkaA’s role in the initial epoxidation, which lead to formation of 3-hydroxy-3-methyloxindole. However, in indole metabolism, indole-2,3-oxide was catalyzed to 3-hydroxyindolin-2-one by a short-chain dehydrogenase, thus further pure enzyme assay should be conducted to explain the formation of 3-hydroxy-3-methyloxindole (Fig. S8) (50, 51). Therefore, this metabolite could originate from two potential routes: successive SkaA-mediated hydroxylation of 3-methyloxindole/3-methyl-3H-indol-3-ol, or enzymatic dehydrogenation (e.g., by a dehydrogenase) of a putative transient skatole dihydrodiol precursor (Fig. S8). The oxygenase of Group E FPMOs is associated with a reductase component, and a reductase gen *skaB* gene was also identified within the cluster (Fig. 2). Although our expression strain did not express the *skaB* gene, it was still capable of catalyzing the skatole reaction. This activity may be attributed to endogenous reductases from *E. coli* that provide the necessary reducing equivalents for SkaA.

Cross-domain similarities are observed in animal CYP450 pathways, supporting the universality of skatole oxidation mechanisms. For example, human CYP450 enzymes could bioactivate skatole via three pathways, i.e., hydroxylation (produce indole-3-carbinol), epoxidation (produce 2,3-epoxide skatole and 3-methyloxindole), and dehydrogenation (produce 3-methyleneindolenine) pathways (52). Pig liver microsomes catalyzed skatole mainly into 3-methyl-3H-indol-3-ol, 3-methyloxidole, and 3-hydroxy-3-methyloxindole (53). What is noteworthy is that the CYP450 gene is widespread in bacterial genomes, suggesting its potential involvement in skatole detoxification and warranting further investigation.

We finally establish SkaA as the founding member of a novel SkaMO subclass within Group E FPMOs (Fig. 5). Detailed analyses of this group of monooxygenases have been extensively conducted by Tischler et al. (30, 54, 55). They proposed that Group E FPMOs can be primarily classified into two types: SMO (which initiates styrene degradation) and IMO (which initiates indole degradation), forming their respective epoxides (30, 39). Our comprehensive phylogenetic reconstruction reveals a distinct third branch characterized by skatole catabolism. Evolutionarily, the parallel emergence of specialized FPMOs for styrene, indole, and skatole—three natural aromatic compounds—suggests substrate-driven divergence. Within Actinobacteria, SkaMOs likely exhibit dual conservation patterns: high sequence conservation (>98% identity) within *R. aetherivorans* and *R. ruber*, versus divergent homologs (∼40% identity) in *Nocardia* that retain genus-level conservation (Tables S8-S9) (20, 24). This implies lineage-specific adaptation to skatole despite shared catalytic mechanisms. Notably, no *Nocardia* skatole degraders have been reported, presenting a research gap. In Gram-negative skatole degraders (*Cupriavidus*, *Pseudomonas*, *Burkholderia*), SkaMO homologs show <40% identity to *Rhodococcus* variants, indicating convergent evolution of distinct molecular solutions (Table S6) (16, 18).

Crucially, multiple strains harbor dual oxygenase systems with partitioned substrate specificity in both Gram-positive (e.g., *R. ruber* R1 and *R. aetherinovorans* BCP1) and Gram-negative strains (e.g., *Burkholderia* sp. IDO3 and *Acinetobacter piscicola* p38) (Table S6) (24, 40, 43). Functional studies confirm this specialization. In *Burkholderia* sp. IDO3, *iifC1* (should encode a SkaMO) was significantly upregulated upon skatole stress, and deletion did not affect indole metabolism, while *iifC2* (should encode an IMO) deletion suppressed indole degradation (40). The gene1687 (XEZ56878, which should encode a SkaMO) in *Acinetobacter piscicola* p38’s was significantly upregulated upon skatole stress, and gene1650 (XEZ56851, which should encode an IMO) was upregulated induced by indole (15). Strikingly, strain p38’s SkaMO could oxidize both skatole and indole, while p38-IMO could only oxide indole. However, strain DMU1 possesses only one *skaA* gene, genetically distinct from those in *R. ruber* R1 and *R. aetherivorans* BCP1. RT-qPCR confirms its involvement in degrading both skatole and indole, indicating complex functional roles based on genomic divergence (Fig. S5). The clear phylogenetic separation of IMO and SkaMO in these strains provides a phylogenetic framework for identifying novel skatole-degrading systems in other microbiota and for predicting the functional roles of different Group E FPMOs (Fig. 5).

## CONCLUSION

This study establishes *Rhodococcus* as a versatile genus capable of widespread skatole degradation across diverse species, revealing previously unrecognized metabolic versatility. We identified a novel skatole epoxidation pathway initiated by the flavin-dependent monooxygenase SkaA. Crucially, phylogenetic analysis redefined Group E FPMOs by establishing SkaMO as a distinct evolutionary clade, exhibiting ≤40% sequence identity to typical styrene/indole mooxygenases and spanning multiple bacterial phyla. The discovery of *skaA*-independent degradation pathways further highlights the genus’s adaptive potential. These findings provide fundamental insights for engineering targeted bioremediation strategies in skatole-polluted environments, and enrich our understanding of the classification Group E FPMOs.

## Acknowledgments

This work was supported by the National Natural Science Foundation of China (No. 32170121).

## Appendix A Supplementary data

Supplementary data associated with this article can be found in the online version.

