## Supplemental materials for "A novel and evolutionarily distinct flavoprotein monooxygenase drives skatole degradation in *Rhodococcus*"

**<u>Supplementary materials for</u>**

| | |
|---|---|
| Tables | 11 |
| Figures | 10 |

Table S1 Primers for this study

| Useage | Gene name | Primer name | Sequence |
|---|---|---|---|
| RT-qPCR | 16S rRNA | DMU1 16S-F | TTCACACATGCTACAATG |
| | | DMU1 16S-R | GCTGATCTGCGATTACTA |
| | DMU1-*skaA* | *skaA*-qF | AGAAGAATCTCCGTGATC |
| | | *skaA*-qR | GTGTAGAGGTCTACTTGG |
| Heterologous expression | pET28a(+) | 28a-his-F | CTCGAGCACCACCACCACC |
| | | 28a-his-R | GGATCCGCGACCCATTTG |
| | DMU1-*skaA* | DMU1-A-F | TCATCACCACAGCCAGGATCCGATGAGAAGAATCTCCGTGAT |
| | | DMU1-A-R | GCATTATGCGGCCGCAAGCTTTCACGCGCTCACCGCCCG |
| | DMU1-*skaB* | DMU1-B-F | TAAGAAGGAGATATACATATGACCACCCACGGACTCGATC |
| | | DMU1-B-R | GGTTTCTTTACCAGACTCGAGCTACCGAGCGGCCGGCGA |
| | SJ-1-*skaA* | SJ-1-His-F | AGCAAATGGGTCGCGGATCCATGACCACCACTCGACGTTCC |
| | | SJ-1-His-R | TGGTGGTGGTGGTGCTCGAGGGCGGGCTGGACCGCTGC |

Table S2 Summary of skatole-degrading strains

| Strain | Degradation ability | Reference |
|---|---|---|
| | **Gram-negative strains** | |
| *Pseudomonas aeruginosa* Gs | Degrade 2 mM skatole within 24 days | (1) |
| *Pseudomonas putida* LPC24 | Degrade 2.0 mM skatole within 30 days | (2) |
| *Acinetobacter* strains NTA1-2A and TAT1-6A | Degrade > 85% skatole (< 200 mg/L) in 6 days | (3) |
| *Acinetobacter oleivorans* AO-06 | Degrade skatole (< 300 mg/L) within 120 h | (4) |
| *Acinetobacter xiamenensis* Ya | Degrade 80 mg/L skatole within 72 h | (5) |
| *Acinetobacter piscicola* p38 | Exploration of potential degradation genes | (6) |
| *Burkholderia* sp. IDO3 | Degrade 50 mg/L skatole within 24 h | (7) |
| Sulfate-reducing bacteria | Degrade 2 mM skatole within 4 weeks | (8) |
| *Cupriavidus* sp. KK10 | Degrade 100 mg/L skatole within 24 h | (9) |
| *Rhodopseudomonas palustris* WKU-KDNS3 | Remove > 90% skatole (0.1 mM) in 21 days | (10) |
| | **Gram-positive strains** | |
| Unidentified Gram-positive bacterium | The removal rate is 32.18% in 4 weeks (100–300 mg/L skatole) | (11) |
| *Lactobacillus brevis* 1.12 | The highest removal rate is 65% in 120 h (1.0 mg/L skatole) | (12) |
| *Rhodococcus aetherivora*ns DMU1 | Degrade 50 mg/L skatole within 24 h | (13) |
| *Rhodococcus pyridinivorans* Rp3 | Degrade 100 mg/L skatole within 48 h | (14) |
| *Rhodococcus ruber* R1 | Degrade 1 mM skatole within 30 h | (15) |
| *Rhodococcus aetherivorans* BCP1 | Degrade 0.5 mM skatole within 48 h | (15) |
| *Bacillus subtilis* GDAAS-A32 | Degrade 100 mg/L skatole within 40 h | (16) |

Table S3 ANI values between the *Rhodococcus* strains

| Strain | DMU1 | DMU2 | DMU114 | DMU2021 | E7 | SJ-1 | SJ-2 | SJ-3 |
|---|---|---|---|---|---|---|---|---|
| **DMU1** | 100 | 72.9 | 78.34 | 77.99 | 72.75 | 91.7 | 76.26 | 75.89 |
| **DMU2** | 72.71 | 100 | 72.31 | 72 | 98.59 | 72.6 | 71.19 | 71.06 |
| **DMU114** | 77.09 | 71.67 | 100 | 97.54 | 71.75 | 77.65 | 77.3 | 76.57 |
| **DMU2021** | 77.03 | 71.63 | 98.11 | 100 | 71.69 | 78.06 | 77.71 | 76.67 |
| **E7** | 72.75 | 98.48 | 72.32 | 72 | 100 | 72.57 | 71.11 | 71.07 |
| **SJ-1** | 90.18 | 72.24 | 77.92 | 78.26 | 72.08 | 100 | 76.44 | 75.13 |
| **SJ-2** | 75.88 | 71.06 | 77.8 | 78.21 | 70.89 | 76.93 | 100 | 98.41 |
| **SJ-3** | 75.72 | 71.01 | 77.18 | 77 | 70.89 | 75.57 | 98.54 | 100 |
| *R. aetherivorans* JCM 14343 | 97.19 | 72.3 | 77.3 | 77.35 | 72.1 | 90.79 | 75.64 | 75.27 |
| *R. antarcticus* 75 | 71.54 | 69.62 | 71.28 | 71.19 | 69.6 | 71.7 | 69.83 | 69.73 |
| *R. artemisiae* YIM 65754 | 76.34 | 71.23 | 77.52 | 77.42 | 71.17 | 76.19 | 87.01 | 87.19 |
| *R. baikonurensis* JCM 11411 | 72.71 | 93.53 | 72.33 | 72.09 | 93.65 | 72.47 | 71.06 | 70.93 |
| *R. cerastii* strain IEGM 1327 | 72.32 | 71.52 | 71.69 | 71.35 | 71.29 | 72.09 | 70.61 | 70.63 |
| *R. cercidiphylli* IEGM 1322 | 72.01 | 71.46 | 71.7 | 71.45 | 71.2 | 72.07 | 70.53 | 70.51 |
| *R. chondri* CC-R104 | 77.7 | 71.72 | 78.16 | 78.24 | 71.59 | 77.55 | 77.38 | 77.94 |
| *R. coprophilus* NCTC10994 | 76.34 | 71.41 | 78.17 | 77.92 | 71.32 | 76.27 | 75.97 | 75.87 |
| *R. daqingensis* CGMCC 1.13630 | 74.73 | 72.75 | 73.80 | 73.56 | 72.72 | 74.72 | 72.3 | 72.15 |
| *R. erythropolis* R138 | 72.63 | 94.88 | 72.25 | 71.86 | 95.15 | 72.51 | 71.01 | 71.02 |
| *R. fascians* DSM 43673 | 71.92 | 71.28 | 71.58 | 71.34 | 71.04 | 71.86 | 70.51 | 70.53 |
| *R. gannanensis* CGMCC 1.15992 | 74.47 | 71.98 | 73.77 | 73.52 | 72.02 | 74.42 | 72.54 | 72.32 |
| *R. globerulus* NBRC 14531 | 72.34 | 79.90 | 71.76 | 71.53 | 79.94 | 72.08 | 70.82 | 70.82 |
| *R. gordoniae* NCTC13296 | 76.95 | 71.5 | 90.01 | 89.87 | 71.55 | 77.01 | 76.64 | 76.55 |
| *R. indonesiensis* SARSHI1 | 90.15 | 72.17 | 77.14 | 77.03 | 72.01 | 94.77 | 75.51 | 75.12 |
| *R. jostii* DSM 44719 | 75.25 | 73.86 | 73.74 | 73.69 | 73.78 | 74.86 | 72.57 | 72.34 |
| *R. koreensis* JCM 10743 | 75.27 | 73.76 | 74 | 73.75 | 73.82 | 75 | 72.52 | 72.29 |
| *R. kroppenstedtii* DSM 44908 | 72.79 | 70.92 | 72.24 | 72.02 | 70.79 | 72.55 | 70.76 | 70.76 |
| *R. kyotonense* JCM 23211 | 72.02 | 71.27 | 71.76 | 71.55 | 71.18 | 72.04 | 70.44 | 70.42 |
| *R. maanshanensis* NBRC 100610 | 74.67 | 72.93 | 73.83 | 73.56 | 72.82 | 74.65 | 72.33 | 72.18 |
| *R. marinonascens* NBRC 14363 | 73.39 | 72.86 | 72.48 | 72.01 | 72.71 | 73.06 | 71.32 | 71.17 |
| *R. navarretei* EXRC-4A-4 | 72.13 | 71.24 | 71.64 | 71.3 | 71.11 | 71.96 | 70.61 | 70.51 |
| *R. olei* JCM 32206 | 75.14 | 72.53 | 74.3 | 73.99 | 72.44 | 75.09 | 72.69 | 72.56 |
| *R. opacus* ATCC 51881 | 75.23 | 73.94 | 73.96 | 73.86 | 73.82 | 74.93 | 72.4 | 72.31 |
| *R. oryzae* NEAU-CX67 | 74.55 | 72.78 | 73.73 | 73.47 | 72.78 | 74.58 | 72.2 | 72.03 |
| *R. oxybenzonivorans* S2-17 | 74.41 | 73.17 | 73.11 | 72.93 | 73.15 | 73.89 | 71.99 | 71.69 |
| *R. parequi* PAM 2766 | 74.92 | 72.74 | 74.28 | 73.99 | 72.71 | 75.06 | 72.75 | 72.72 |
| *R. pseudokoreensis* R79 | 75.41 | 73.77 | 74.09 | 73.89 | 73.84 | 75.08 | 72.58 | 72.33 |
| *R. pyridinivorans* DSM 44555 | 77.09 | 71.65 | 97.92 | 98.4 | 71.72 | 77.31 | 77.02 | 76.66 |

| | | | | | | | | |
|---|---|---|---|---|---|---|---|---|
| *R. qingshengii* JCM 15477 | 72.64 | 98.35 | 72.41 | 71.93 | 98.5 | 72.52 | 71.07 | 70.99 |
| *R. rhodnii* ATCC 35071 | 73.31 | 71.26 | 72.95 | 72.71 | 71.24 | 73.13 | 71.44 | 71.47 |
| *R. rhodochrous* NCTC10210 | 77.01 | 71.68 | 93.77 | 93.24 | 71.56 | 77.14 | 76.71 | 76.54 |
| *R. ruber* DSM 43338 | 90.29 | 72.19 | 77.3 | 77.05 | 72.13 | 98.76 | 75.25 | 75.11 |
| *R. sacchari* Z13 | 77.52 | 71.78 | 82.98 | 82.74 | 71.82 | 77.39 | 76.45 | 76.33 |
| *R. sovatensis* 18930 | 71.48 | 70.79 | 71.02 | 70.78 | 70.8 | 71.33 | 69.9 | 69.95 |
| *R. spelaei* C9-5 | 74.73 | 72.98 | 73.96 | 73.69 | 72.97 | 74.72 | 72.43 | 72.32 |
| *R. spongiicola* LHW50502 | 74.19 | 72.23 | 73.37 | 73.24 | 72.18 | 74.01 | 71.96 | 72.11 |
| *R. triatomae* DSM 44893 | 74.13 | 72.33 | 73.53 | 73.19 | 72.26 | 74.01 | 71.97 | 71.96 |
| *R. trifolii* CCM 7905 | 71.66 | 70.76 | 71.3 | 71 | 70.66 | 71.59 | 70.14 | 70.19 |
| *R. tukisamuensis* JCM 11308 | 74.13 | 72.33 | 73.53 | 73.19 | 72.26 | 74.01 | 71.97 | 71.96 |
| *R. wratislaviensis* YZ02 | 75.37 | 73.93 | 73.86 | 73.69 | 73.86 | 74.95 | 72.44 | 72.24 |
| *R. xishaensis* LHW51113 | 74.2 | 72.17 | 73.42 | 73.22 | 72.02 | 73.89 | 72 | 72.05 |
| *R. yananensis* FBM22-1 | 76.74 | 71.47 | 77.57 | 77.29 | 71.51 | 76.69 | 77.35 | 77.25 |
| *R. yunnanensis* NBRC 103083 | 71.71 | 71.03 | 71.4 | 71.1 | 70.99 | 71.67 | 70.16 | 70.19 |
| *R. zopfii* NBRC 100606 | 78.75 | 71.97 | 77.66 | 77.97 | 71.88 | 78.49 | 76.23 | 76.07 |

Table S4 dDDH values between the *Rhodococcus* strains

| Query strain | Subject strain | dDDH (d4, in %) | G+C content difference (in %) |
|---|---|---|---|
| DMU1 | *R. aetherivorans* JCM 14343 | 91.4 | 0.24 |
| DMU1 | *R. aetherivorans* DSM 44752 | 91.3 | 0.23 |
| DMU2 | E7 | 89.9 | 0 |
| DMU2 | *R. jialingiae* djl-6-2 | 89.1 | 0.08 |
| DMU2 | *R. qingshengii* JCM 15477 | 88.5 | 0.13 |
| DMU2 | *R. enclensis* DSM 45688 | 87.9 | 0.19 |
| DMU114 | *R. pyridinivorans* DSM 44555 | 85.7 | 0.03 |
| DMU114 | DMU2021 | 86.2 | 0.22 |
| DMU2021 | *R. pyridinivorans* DSM 44555 | 91.1 | 0.19 |
| DMU2021 | *R. biphenylivorans* TG9 | 88.4 | 0.39 |
| E7 | *R. enclensis* DSM 45688 | 89.9 | 0.19 |
| E7 | *R. qingshengii* JCM 15477 | 89.5 | 0.13 |
| SJ-1 | *R. ruber* NBRC 15591 | 92.9 | 0.32 |
| SJ-1 | *R. ruber* DSM 43338 | 92.7 | 0.3 |
| SJ-2 | SJ-3 | 91.6 | 0.02 |
| SJ-2 | *R. artemisiae* YIM 65754 | 33.9 | 0.24 |
| SJ-3 | *R. artemisiae* YIM 65754 | 34.1 | 0.22 |

Table S5 Potential SkaA homologs (>30% similarity) in all the 53 *Rhodococcus* strains from table S4

| Strain | Subject ID | Percent identity (%) | Query coverage | E-value |
|---|---|---|---|---|
| *Rhodococcus aetherivorans* DMU1 | QIX49842 | 100 | - | - |
| *Rhodococcus aetherivorans* JCM 14343 | WP_152103792.1 | 99.3 | 100 | 0 |
| *Rhodococcus aetherivorans* JCM 14343 | WP_043796862.1 | 38.1 | 80.7 | 3.35e-67 |
| *Rhodococcus indonesiensis* SARSHI1 | WP_416062207.1 | 38.4 | 80.7 | 2.82e-69 |
| *Rhodococcus opacus* ATCC 51881 | WP_005563102.1 | 37.9 | 80.7 | 6.01e-65 |
| *Rhodococcus opacus* ATCC 51881 | WP_005568880.1 | 33.4 | 91.4 | 2.73e-65 |
| *Rhodococcus opacu*s ATCC 51881 | WP_005568881.1 | 30.7 | 91.4 | 4.68e-53 |
| *Rhodococcus oxybenzonivorans* S2-17 | WP_109335618.1 | 30.0 | 92.1 | 1.17e-30 |
| *Rhodococcus ruber* SJ-1 | WP_010594217.1 | 38.2 | 80.7 | 1.23e-66 |
| *Rhodococcus sacchari* Z13 | WP_264154094.1 | 36.9 | 92.3 | 3.32e-65 |
| *Rhodococcus spongiicola* LHW 50502 | WP_127947546.1 | 34.0 | 91.4 | 5.20e-65 |
| *Rhodococcus spongiicola* LHW 50502 | WP_127947545.1 | 31.9 | 91.7 | 3.31e-55 |
| *Rhodococcus tukisamuensis* JCM 11308 | WP_072736178.1 | 36.6 | 80.7 | 1.57e-60 |

Table S6 Distribution of potential SkaA homologs (>30% similarity) in all available genomes of skatole-degrading strains

| **Strain** | **Protein ID** | **Start** | **End** | **Gene length** | **Similarity** |
|---|---|---|---|---|---|
| *Rhodococcus aetherivorans* DMU1 | QIX49842 | 2091562 | 2092929 | 1368 | 100% |
| *Rhodococcus ruber* SJ-1 | WP_010594217 | 3286923 | 3288296 | 1374 | 38.2% |
| *Rhodococcus ruber* R1 | WP_161572325 | 1670408 | 1671775 | 1368 | 98.9% |
| *Rhodococcus ruber* R1 | WP_010594217 | 5180175 | 5181548 | 1374 | 38.2% |
| *Rhodococcus aetherinovorans* BCP1 | WP_155951766 | 1292779 | 1294146 | 1368 | 99.1% |
| *Rhodococcus aetherinovorans* BCP1 | WP_006943660 | 5068421 | 5069794 | 1374 | 38.2% |
| *Cupriavidus* sp. KK10 | WP_008642920 | 6373548 | 6374789 | 1242 | 34.8% |
| *Cupriavidus* sp. KK10 | WP_013953986 | 3671115 | 3672404 | 1290 | 32.8% |
| *Burkholderia* sp. IDO3 | APT36898 | 820804 | 822069 | 1266 | 35.7% |
| *Burkholderia* sp. IDO3 | AXK62974 | 2226214 | 2227458 | 1245 | 32.2% |
| *Acinetobacter piscicola* p38 | XEZ56851 | 1690588 | 1691826 | 1239 | 31.6% |
| *Acinetobacter piscicola* p38 | XEZ56878 | 1723205 | 1724476 | 1272 | 36.9% |
| *Rhodococcus qingshengii* DMU2 | - | - | - | - | - |
| *Rhodococcus pyridinivorans* DMU114 | - | - | - | - | - |
| *Rhodococcus pyridinivorans* DMU2021 | - | - | - | - | - |
| *Rhodococcus qingshengii* E7 | - | - | - | - | - |
| *Rhodococcus* sp. SJ-2 | - | - | - | - | - |
| *Rhodococcus* sp. SJ-3 | - | - | - | - | - |
| *Rhodococcus pyridinivorans* Rp3 | - | - | - | - | - |

Table S7 The gene cluster responsible for skatole degradation in strain DMU1

| Gene | Protein ID | Name | Homologous protein |
|---|---|---|---|
| HFP48_09850 | QIX49835.1 | SkaC | Aromatic-ring-hydroxylating dioxygenase subunit alpha |
| HFP48_09855 | QIX49836.1 | SkaD | Aromatic-ring-hydroxylating dioxygenase subunit beta |
| HFP48_09860 | QIX49837.1 | SkaZ | Unknow |
| HFP48_09865 | QIX49838.1 | SkaE | Type 1 glutamine amidotransferase |
| HFP48_09870 | QIX49839.1 | SkaF | NAD(P)/FAD-dependent oxidoreductase |
| HFP48_09875 | QIX49840.1 | SkaG | 4Fe-4S binding protein |
| HFP48_09880 | QIX49841.1 | SkaH | Glutamine synthetase |
| HFP48_09885 | QIX49842.1 | SkaA | Styrene monooxygenase StyA |
| HFP48_09890 | QIX49843.1 | SkaI | Ectoine hydrolase |
| HFP48_09895 | QIX49843.1 | SkaJ | Hydantoinase subunit B |
| HFP48_09900 | QIX49845.1 | SkaK | Hydantoinase subunit A |
| HFP48_09905 | QIX49846.1 | SkaR | LuxR C-terminal-related transcriptional regulator |
| HFP48_09910 | QIX49847.1 | SkaL | SDR family oxidoreductase |
| HFP48_09915 | QIX49848.1 | SkaB | Flavin reductase family protein |

Table S8 SkaA homologs in the NCBI RefSeq database with ≥40% identity

| ID | Similarity | Genus | Species | Strain |
|---|---|---|---|---|
| WP_149433161.1 | 40.7 | *Antrihabitans* | *cavernicola* | |
| WP_237821021.1 | 58.7 | *Arthrobacter* | *hankyongi* | |
| WP_283501880.1 | 63.3 | *Cryobacterium* | sp. | PH31-O1 |
| WP_283497437.1 | 62.4 | *Cryobacterium* | sp. | PH29-G1 |
| WP_134502114.1 | 63.7 | *Cryobacterium* | *glaciale* | |
| WP_170194208.1 | 42.3 | *Gordonia* | *asplenii* | |
| WP_045119590.1 | 50.5 | *Haliangium* | *ochraceum* | |
| WP_402645705.1 | 40.9 | *Nocardia* | sp. | NPDC 087230 |
| WP_396916410.1 | 40.6 | *Nocardia* | *brasiliensis* | |
| WP_393442621.1 | 40.1 | *Nocardia* | sp. | NPDC 049149 |
| WP_387700286.1 | 40.9 | *Nocardia* | *thailandica* | |
| WP_379315658.1 | 40.9 | *Nocardia* | sp. | NPDC 060220 |
| WP_357240208.1 | 40.9 | *Nocardia* | sp. | NPDC 019395 |
| WP_345494243.1 | 42 | *Nocardia* | *callitridis* | |
| WP_342214927.1 | 41.3 | *Nocardia* | *cyriacigeorgica* | |
| WP_280441622.1 | 40.6 | *Nocardia* | *brasiliensis* | |
| WP_280430607.1 | 40.3 | *Nocardia* | *brasiliensis* | |
| WP_280409824.1 | 40.9 | *Nocardia* | *brasiliensis* | |
| WP_280404053.1 | 40.6 | *Nocardia* | *brasiliensis* | |
| WP_280393585.1 | 40.6 | *Nocardia* | *brasiliensis* | |
| WP_195137510.1 | 40.6 | *Nocardia* | *brasiliensis* | |
| WP_195091645.1 | 40.6 | *Nocardia* | *brasiliensis* | |
| WP_043663825.1 | 40.6 | *Nocardia* | *vulneris* | |
| WP_043651352.1 | 41.1 | *Nocardia* | *thailandica* | |
| WP_042258574.1 | 40.3 | *Nocardia* | *brasiliensis* | |
| WP_029900882.1 | 40.6 | *Nocardia* | *brasiliensis* | |
| WP_014988896.1 | 40.6 | *Nocardia* | *brasiliensis* | |
| WP_252568485.1 | 58.3 | *Pseudarthrobacter* | sp. | HLT3-5 |
| WP_263507466.1 | 99.3 | *Rhodococcus* | *aetherivorans* | |
| WP_203329394.1 | 99.6 | *Rhodococcus* | | |
| WP_185113676.1 | 99.8 | *Rhodococcus* | *aetherivorans* | |
| WP_170914061.1 | 99.6 | *Rhodococcus* | | |
| WP_168579423.1 | 100 | *Rhodococcus* | *aetherivorans* | DMU1 |
| WP_161995899.1 | 99.6 | *Rhodococcus* | sp. | YH1 |
| WP_161572325.1 | 98.9 | *Rhodococcus* | *ruber* | |
| WP_155951766.1 | 99.1 | *Rhodococcus* | *aetherivorans* | |
| WP_152103792.1 | 99.3 | *Rhodococcus* | *aetherivorans* | |
| WP_146034277.1 | 99.6 | *Rhodococcus* | | |
| WP_146560306.1 | 40 | *Tsukamurella* | *asaccharolytica* | |
| WP_108076155.1 | 40.1 | *Vitiosangium* | sp. | GDMCC 1.1324 |

Table S9 SkaA homologs in the NCBI NR database with ≥40% identity

| ID | Similarity | Genus | Species | Strain |
|---|---|---|---|---|
| WP_283501880.1 | 63.3 | *Cryobacterium* | sp. | PH |
| WP_283497437.1 | 62.4 | *Cryobacterium* | sp. | PH |
| WP_280441622.1 | 40.6 | *Nocardia* | *brasiliensis* | |
| WP_280430607.1 | 40.3 | *Nocardia* | *brasiliensis* | |
| WP_280409824.1 | 40.9 | *Nocardia* | *brasiliensis* | |
| WP_280404053.1 | 40.6 | *Nocardia* | *brasiliensis* | |
| WP_280393585.1 | 40.6 | *Nocardia* | *brasiliensis* | |
| WP_280230640.1 | 40.4 | *Nocardia* | *cyriacigeorgica* | |
| WP_280206591.1 | 40.2 | *Nocardia* | *cyriacigeorgica* | |
| WP_263507466.1 | 99.3 | *Rhodococcus* | *aetherivorans* | |
| WP_252568485.1 | 58.3 | *Pseudarthrobacter* | sp. | HLT |
| WP_237821021.1 | 58.7 | *Arthrobacter* | *hankyongi* | |
| WP_203329394.1 | 99.6 | *Rhodococcus* | | |
| WP_195137510.1 | 40.6 | *Nocardia* | *brasiliensis* | |
| WP_195091645.1 | 40.6 | *Nocardia* | *brasiliensis* | |
| WP_185113676.1 | 99.8 | *Rhodococcus* | *aetherivorans* | |
| WP_170914061.1 | 99.6 | *Rhodococcus* | sp. | M |
| WP_170194208.1 | 42.3 | *Gordonia* | *asplenii* | |
| WP_168579423.1 | 100 | *Rhodococcus* | sp. | DMU1 |
| WP_161572325.1 | 98.9 | *Rhodococcus* | *ruber* | |
| WP_155951766.1 | 99.1 | *Rhodococcus* | *aetherivorans* | |
| WP_152103792.1 | 99.3 | *Rhodococcus* | *aetherivorans* | |
| WP_149433161.1 | 40.7 | *Spelaeibacter* | *cavernicola* | |
| WP_146560306.1 | 40 | *Tsukamurella* | *asaccharolytica* | |
| WP_146034277.1 | 99.6 | *Rhodococcus* | *aetherivorans* | |
| WP_134502114.1 | 63.7 | *Cryobacterium* | *glaciale* | |
| WP_108076155.1 | 40.1 | *Vitiosangium* | sp. | GDMCC |
| WP_045119590.1 | 50.5 | *Haliangium* | *ochraceum* | |
| WP_043663825.1 | 40.6 | *Nocardia* | *vulneris* | |
| WP_043651352.1 | 41.1 | *Nocardia* | *thailandica* | |
| WP_042258574.1 | 40.3 | *Nocardia* | *brasiliensis* | |
| WP_029900882.1 | 40.6 | *Nocardia* | *brasiliensis* | |
| WP_014988896.1 | 40.6 | *Nocardia* | *brasiliensis* | |
| OLL18304.1 | 99.3 | *Rhodococcus* | sp. | M8 |
| NCL75270.1 | 99.6 | *Rhodococcus* | sp. | YH |
| MDQ1502227.1 | 57.1 | unknown | bacterium | |
| MCU1347542.1 | 41.8 | unknown | bacterium | |
| MCI0543486.1 | 52.6 | unknown | bacterium | |
| MCI0425126.1 | 52.3 | unknown | bacterium | |
| MCI0408171.1 | 40.9 | unknown | bacterium | |
| MCH8128790.1 | 47.3 | unknown | bacterium | |

| | | | | |
|---|---|---|---|---|
| MCH8056365.1 | 40.8 | unknown | bacterium | |
| MCH7845050.1 | 46.7 | unknown | bacterium | |
| KDE14434.1 | 98.9 | *Rhodococcus* | *aetherivorans* | |
| HYV43985.1 | 40.2 | *Myxococcaceae* | bacterium | |
| HWN71841.1 | 46.9 | *Haliangium* | sp | |
| HVV87497.1 | 41 | *Kofleriaceae* | bacterium | |
| HVR78000.1 | 47.4 | *Acidimicrobiia* | bacterium | |
| HVE81196.1 | 40.2 | *Myxococcales* | bacterium | |
| HUF96357.1 | 48.8 | *Acidimicrobiia* | bacterium | |
| HMV69958.1 | 44.1 | unknown | bacterium | |
| HEX2573964.1 | 40 | *Polyangia* | bacterium | |
| GES35234.1 | 99.1 | *Rhodococcus* | *aetherivorans* | |
| CCW14454.1 | 99.6 | *Rhodococcus* | *aetherivorans* | |
| ASF11060.2 | 40.2 | *Nocardia* | *brasiliensis* | |
| ANZ25892.1 | 99.3 | *Rhodococcus* | sp. | WB1 |
| ACY18693.1 | 50.8 | *Haliangium* | *ochraceum* | |

Table S10 Flavoprotein monooxygenases used in this study

| Group | Annotation | Name | Resource strain |
|---|---|---|---|
| Group A | 6-Hydroxynicotinate 3-monooxygenase 6HNMO | PDB 5EOW | *Pseudomonas putida* KT2440 |
| Group A | p-Nitrophenol 4-monooxygenase PnpA | 6AIN_A | *Pseudomonas putida* |
| Group A | FAD-dependent monooxygenase tropB | 6NEV_A | *Talaromyces stipitatus* ATCC 10500 |
| Group B | Phenylacetone monooxygenase | 4D04_A | *Thermobifida fusca* |
| Group B | Cyclohexanone monooxygenase | 4RG4_A | *Rhodococcus* sp. HI-31 |
| Group B | Steroid monooxygenase | 4AOS_A | Rhodococcus *rhodochrous* |
| Group C | Dibenzothiophene-sulfone monooxygenase | Q0ZIH7 | *Rhodococcus erythropolis* |
| Group C | Nitrilotriacetate monooxygenase | 3SDO_A | *Burkholderia pseudomallei* |
| Group C | Long-chain alkane monooxygenase | A4IU28 | *Geobacillus thermodenitrificans* |
| Group D | 4-Nitrophenol monooxygenase | UniProtKB Q6F4M8 | *Rhodococcus opacus* |
| Group D | 4-Hydroxyphenylacetate 3-monooxygenase oxygenase | 6QYI_A | *Escherichia coli* |
| Group D | Chlorophenol-4-monooxygenase | 4OO2_A | *Streptomyces globisporus* |
| Group E | Styrene monooxygenase | UniProtKB O50214 | *Pseudomonas* sp. VLB120 |
| Group E | Styrene monooxygenase | O06834 | *Pseudomonas fluorescens* ST |
| Group E | Indole monooxygenase IifC | ARO76328.1 | *Acinetobacter* sp. O153 |
| Group E | Indole monooxygenase IifC | AXK62974 | *Burkholderia* sp. IDO3 |
| Group F | Tryptophan 7-halogenase KtzQ | A8CF75 | *Kutzneria* sp. 744 |
| Group F | Tryptophan halogenase | 6FRL_A | *Brevundimonas* sp. BAL3. |
| Group F | Tryptophan 5-halogenase (PyrH) | 2WEU_A | *Streptomyces rugosporus* |
| Group G | Lysine 2-monooxygenase LMO | BAG54787 | *Pseudomonas putida* |
| Group G | L-phenylalanine oxidase | Q5W9R9 | *Pseudomonas* sp. P-501 |

| | | | |
|---|---|---|---|
| Group G | Tryptophan 2-monooxygenase | 4IV9_A | *Pseudomonas savastanoi* |
| Group H | Lactate 2-monooxygenase | AMO62034 | *Mycolicibacterium phle*i |
| Group H | Nitronate monooxygenase | AVR66134 | *Pseudomonas paraeruginosa* |
| Group H | Nitronate monooxygenase | 6BKA_A | *Cyberlindnera mrakii* |

Table S11 Comparison of SkaA with reported styrene and indole monooxygenases. The colors in the table match those in Figure 5.

| **Description** | **Identity (%)** |
|---|---|
| WP_168579423.1 SkaA *Rhodococcus* sp. DMU1 | 100 |
| WP_155951766.1 SkaA *Rhodococcus aetherivorans* BCP1 | 99.1 |
| WP_161572325.1 SkaA *Rhodococcus ruber* R1 | 98.9 |
| APT36898.1 IifC1 *Burkholderia* sp. IDO3 | 35.7 |
| ASR05096.1 GrIndA *Gordonia rubripertincta* CWB2 | 35.7 |
| WP_010594217.1 SkaA *Rhodococcus ruber* SJ-1 | 38.2 |
| XEZ56878.1 SkaA *Acinetobacter piscicola* p38 | 36.9 |
| WP_027965539.1 HhIndA *Halomonas halocynthiae* DSM14573 | 34.5 |
| CAJ94554.1 CnIndA *Cupriavidus necator* H16 | 35.0 |
| ADU39062.1 VpIndA2B *Variovorax paradoxus* EPS | 34.2 |
| WP_028028710.1 GnIndA *Gemmobacter nectariphilus* 15620 | 32.6 |
| EEA04565.1 BH160IndA *Burkholderia* sp. H160 | 33.0 |
| EJJ03822.1 SaIndA1 *Streptomyces auratus* AGR0001 | 35.1 |
| WP_039783212.1 HhMO *Herbaspirillum huttiense* subsp. *putei* IAM 15032 | 31.9 |
| ACR43973.1 RoIndA1 *Rhodococcus opacus* 1CP | 33.3 |
| KWR77134.1 CSHEIndA *Cupriavidus* sp. SHE | 33.3 |
| AXK62974.1 IifC2 *Burkholderia* sp. IDO3 | 32.2 |
| BAD56093.1 NfIndA1 *Nocardia farcinica* IFM 10152 | 33.5 |
| ADU39063.1 VpIndA1 *Variovorax paradoxus* EPS | 34.7 |
| ABM07034.1 PaIndA1 *Paenarthrobacter aurescens* TC1 | 32.3 |
| XEZ56851.1 IndA *Acinetobacter piscicola* p38 | 31.6 |
| ARO76328.1 IifC *Acinetobacter* sp. O153 | 30.4 |
| ENW75087.1 IifC *Acinetobacter baumannii* ATCC 19606 | 31.6 |
| WP_081399512.1 IifC *Acinetobacter* sp. JW | 30.5 |
| CAG69430.1 AbIndA *Acinetobacter baylyi* ADP1 | 30.4 |
| ACR43974.1 RoIndA2B *Rhodococcus opacus* 1CP | 32.1 |
| ABQ12175.1 MoxY uncultured bacterium | 32.8 |
| ABX34433.1 DaIndA *Delftia acidovorans* SPH-1 | 32.3 |
| EJJ03821.1 SaIndA2B *Streptomyces auratus* AGR0001 | 31.6 |
| BAD56094.1 NfIndA2B *Nocardia farcinica* IFM 10152 | 32.2 |
| ABM10099.1 PaIndA2B *Paenarthrobacter aurescens* TC1 | 30.3 |
| ABZ79366.1 MoxA uncultured bacterium | 27.7 |
| ABH00103.1 RjStyA *Rhodococcus jostii* RHA1 | 29.4 |
| WP_027934242.1 AtStyA *Amycolatopsis thermoflava* N1165 | 27.7 |
| WP_019810476.1 ShStyA *Saccharomonospora halophila* 8 | 29.6 |
| ABH00101.1 RjStyA *Rhodococcus jostii* RHA1 | 27.4 |
| WP_091504755.1 AsMO *Amycolatopsis sacchari* DSM 44468 | 28.0 |
| WP_030740546.1 StrepStyA *Streptomyces* sp. NRRL S-31 | 27.3 |
| ABV24041.1 SmoA uncultured | 25.6 |
| GAB22407.1 GpStyA *Gordonia polyisoprenivorans* NBRC 16320 | 25.4 |

| | |
|---|---|
| BAL04132.1 RoST5StyA *Rhodococcus* sp. ST-5 | 29.0 |
| BAL04129.1RoST10StyA *Rhodococcus* sp. ST-10 | 27.3 |
| ASR05591.1 GrStyA *Gordonia rubripertincta* CWB2 | 28.7 |
| WP_022977994.1 NrStyA *Nevskia ramosa* DSM11499 | 26.7 |
| GAC06215.1 PaStyA *Paraglaciecola agarilytica* NO2 | 25.7 |
| WP_039579272.1 SfStyA *Sphingopyxis fribergensis* Kp5.2 | 27.3 |
| ANS32444.1 RoStyA *Rhodococcus opacus* 1CP | 27.8 |
| WP_027855270.1 MlStyA *Marinobacterium litorale* DSM 23545 | 24.4 |
| CAA04000.1 PY2StyA *Pseudomonas* sp. Y2 | 24.0 |
| ADE62390.1 PLQ26StyA *Pseudomonas* sp. LQ26 | 25.3 |
| CAA73790.1 PpStyA S12 *Pseudomonas putida* S12 | 24.1 |
| AAC23718.1 PtStyA *Pseudomonas* sp. VLB120 | 25.1 |
| ABX24519.1 PpStyA CA-3 *Pseudomonas putida* CA-3 | 24.1 |
| CAB06823.1 PfStyA *Pseudomonas fluorescens* ST | 23.8 |
| ABB03727.1 PpStyA SN1 *Pseudomonas putida* SN1 | 24.0 |

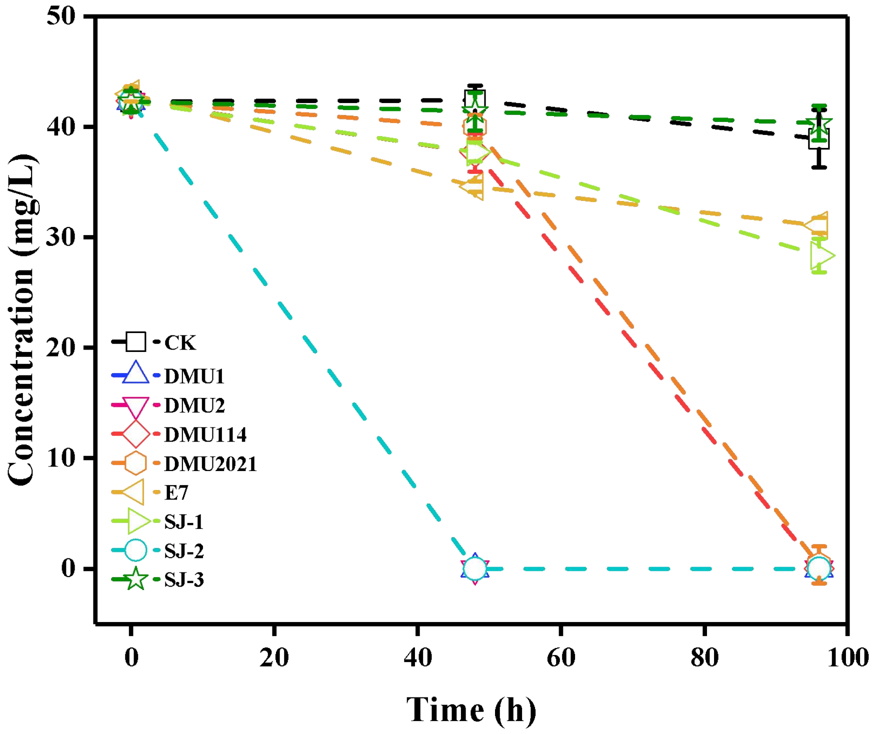


Fig. S1 Skatole-degrading curves of eight *Rhodococcus* strains in MS medium with skatole as the sole carbon source

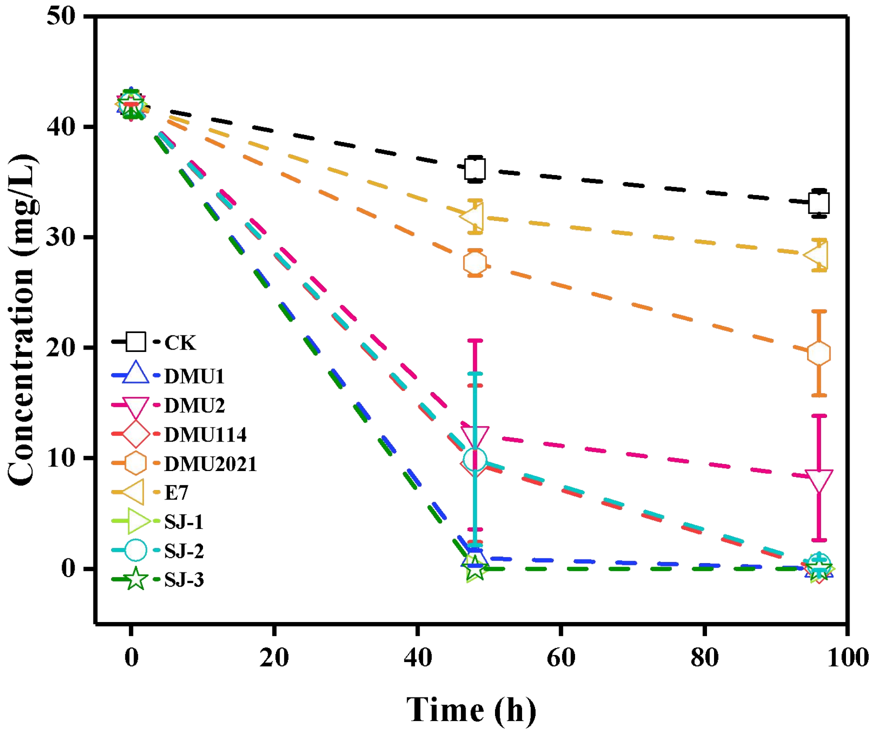


Fig. S2 Skatole-degrading curves of eight *Rhodococcus* strains in MSY medium with yeast extract as the extra carbon source

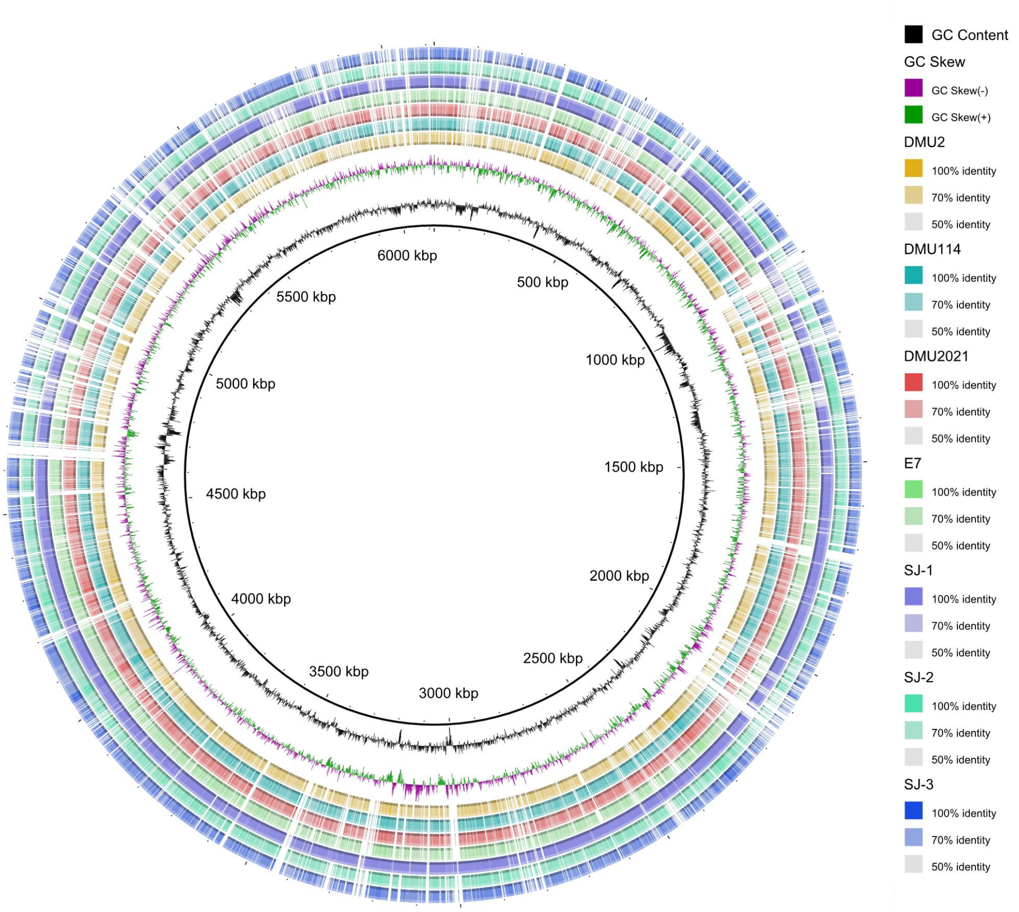


Fig. S3 Comparative genomic circular diagram of the eight *Rhodococcus* strains

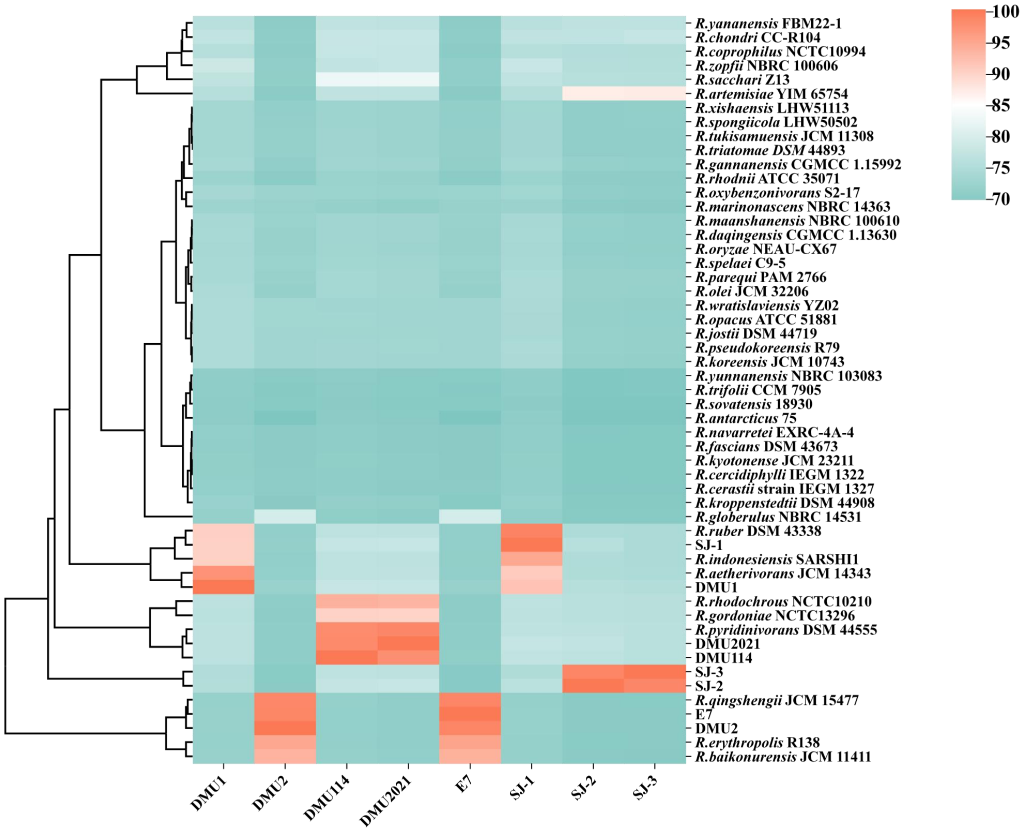


Fig. S4 Average nucleotide identity (ANI) analysis of eight strains with the genomes of valid *Rhodococcus* species.

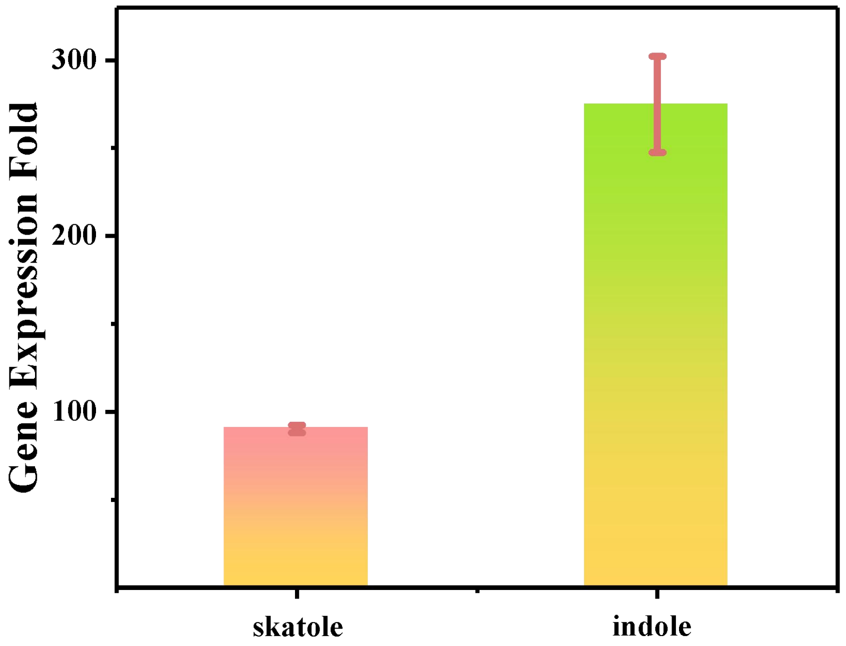


Fig. S5 RT-qPCR results of gene *skaA* in strain DMU1 in response to skatole and indole.

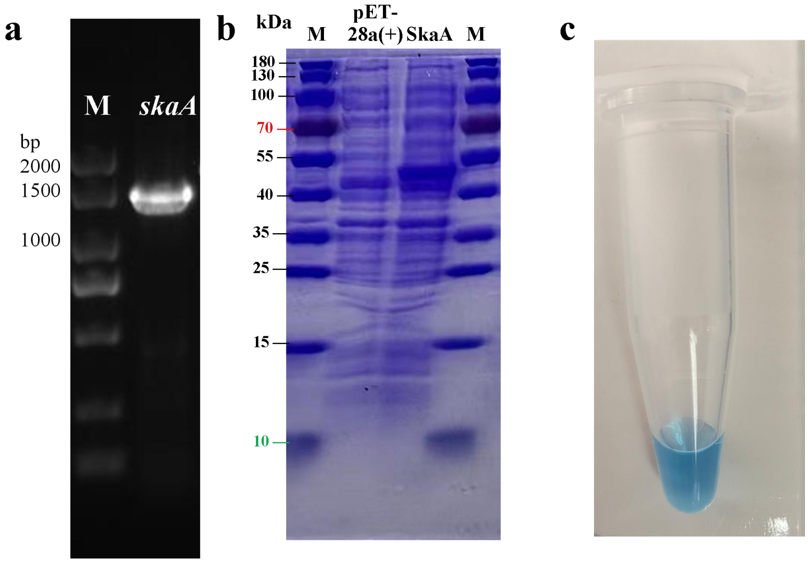


Fig. S6 Gene amplification and expression results. (a) PCR products of gene *skaA*. (b) SDS-PAGE of *E. coli* BL21(DE3) expressing *skaA* gene. (c)Indigoids formation with indole as the substrate.

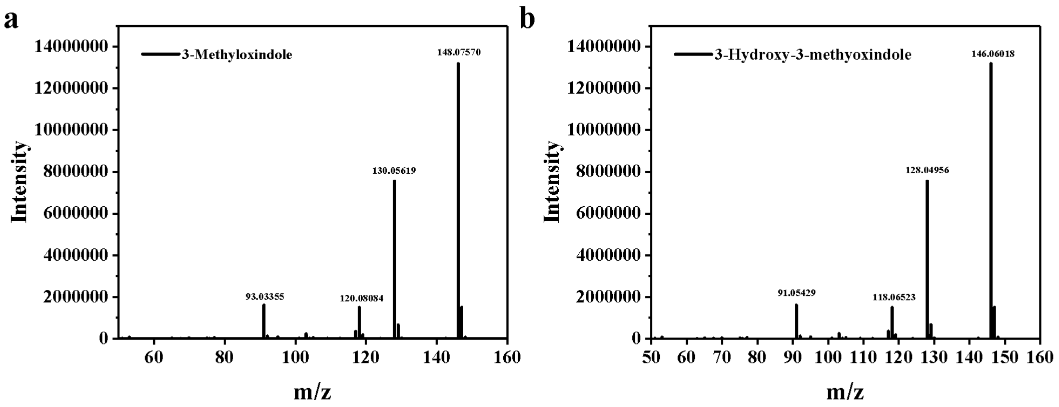


Fig. S7 Mass spectra of the standard compounds of (a) 3-methyloxindole and (b) 3-hydroxy-3-methyloxindole.

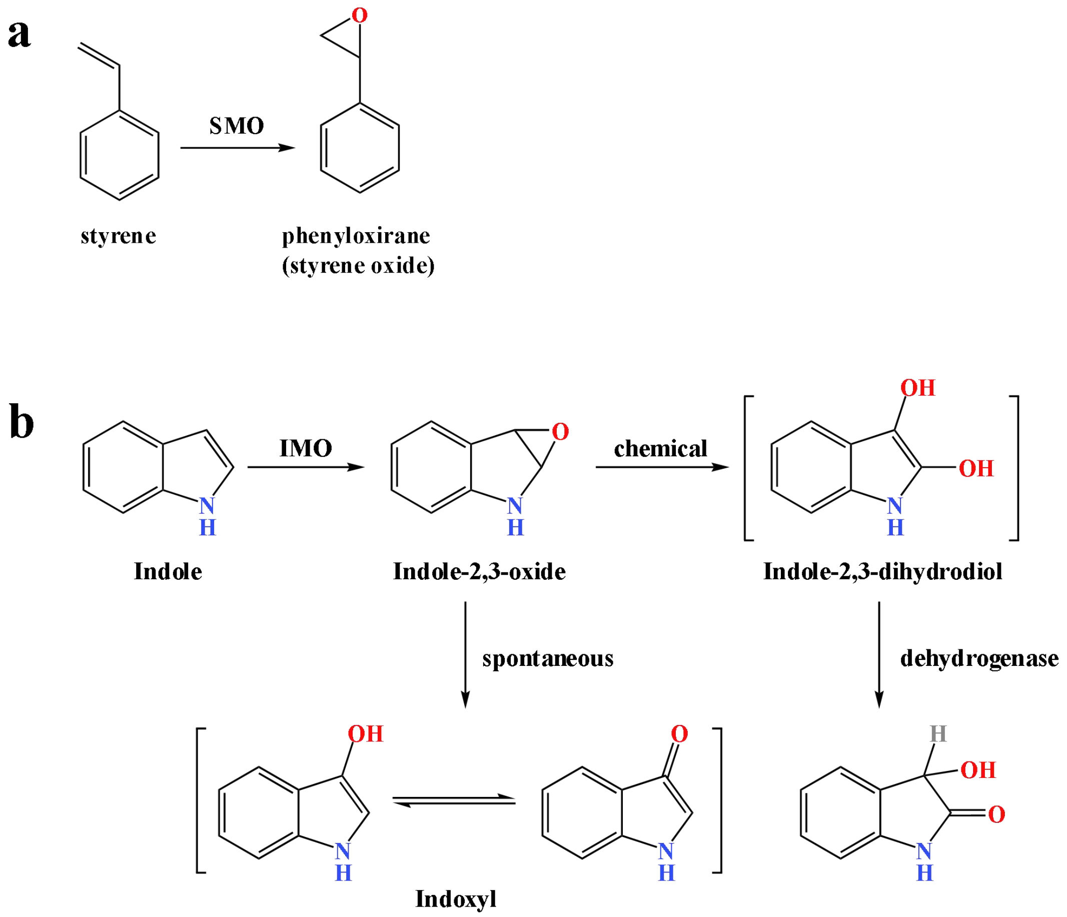


Fig. S8 Typical reactions catalyzed by (a) styrene monooxygenase (SMO) and (b) indole monooxygenase (IMO).

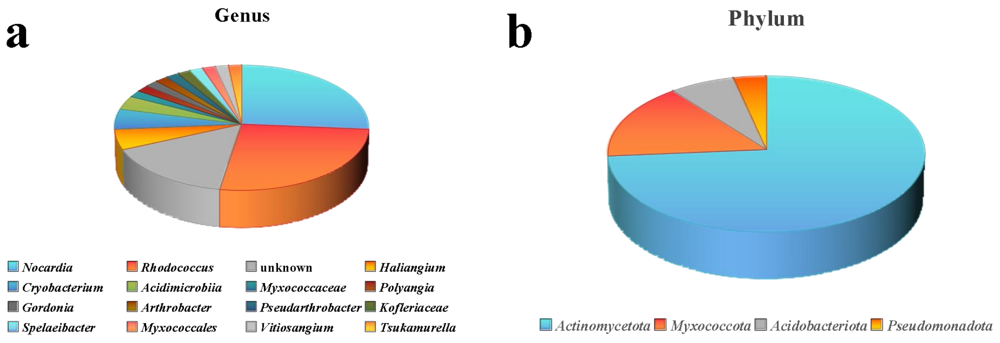


Fig. S9 Distribution of SkaA homologs in related strains in the NCBI non-redundant (NR) database. (a) Genus. (b) Phylum.

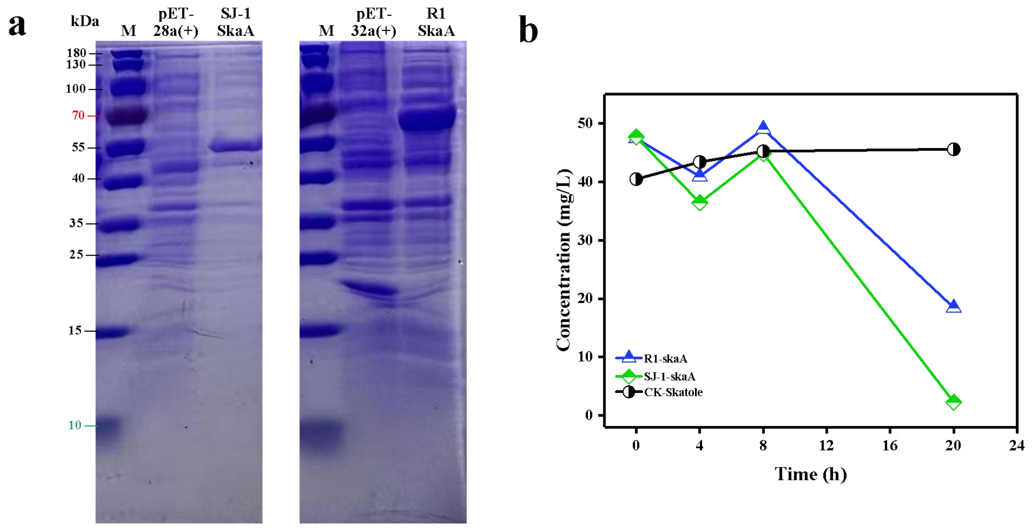


Fig. S10 Gene expression and skatole degradation performance of SkaA homologs. (a) Expression of SkaA homologs in strains SJ-1 and R1. (b) Skatole degradation performance of the corresponding gene-engineered strains.
